# One-pot chemo-enzymatic synthesis and one-step recovery of homogeneous long-chain polyphosphates from microalgal biomass

**DOI:** 10.1101/2023.08.19.553819

**Authors:** Yi-Hsuan Lin, Shota Nishikawa, Tony Z. Jia, Fang-I Yeh, Anna Khusnutdinova, Alexander F. Yakunin, Kosuke Fujishima, Po-Hsiang Wang

## Abstract

Graphical abstract

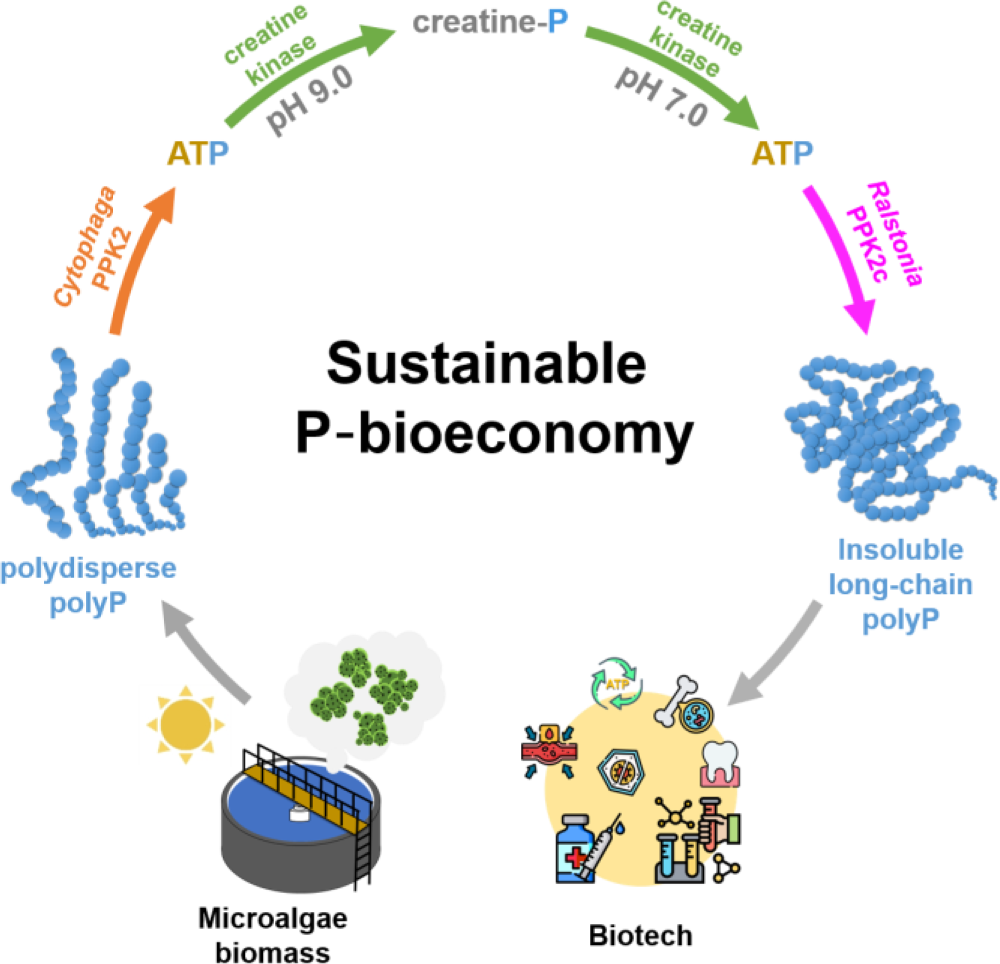

Phosphate, an essential component of life, fertilizers, and detergents, is a finite resource that could be depleted within 70 years, while improper phosphate waste disposal in aquatic environments results in eutrophication. Despite some chemical-based methods, biological phosphorus removal using polyphosphate-accumulating organisms, such as microalgae, serves as a sustainable alternative to reclaim phosphate from wastewater. Polyphosphates have profound biological functions and biomedical applications, serving as energy stock, coagulation factors, and antiviral agents depending on their length, showing inherent value in polyphosphate recovery. Here, we leveraged the power of thermodynamic coupling and phase transitions to establish a one-pot, two-step multi-enzyme cascade to convert polydisperse polyphosphate in microalgae biomass into high-molecular-weight insoluble long-chain polyphosphates, allowing for one-step purification. We then optimzed a thermo-digestion approach to transform the 1,300-mers into shorter polyphosphates. Altogether, the processes established here enable the establishment of a sustainable P bioeconomy platform to refine microalgal biomass for biotechnological uses.

## Introduction

Phosphorus is a key element in the biomass of all living organisms ^1^ and is also essential for modern agriculture and industry as a component in fertilizer, animal feed, and detergents ^2^. However, most of the accessible phosphorus sources exist in the form of apatite minerals in the lithosphere and are inaccessible to land-based plants, while worldwide phosphorus demand has been rapidly growing and is expected to exceed supply within 70 years due to the rapid increase in global population ^3^. To increase the phosphorus supply, “wet process methods” have been invented to convert unusable inorganic phosphorus into phosphoric acid, a precursor to fertilizers, followed by an introduction to land plants ^4^. However, the excessive introduction of soluble phosphorus into the aquatic environments also causes detrimental impacts ^5^, *e.g.*, phosphorus leakage from agricultural fields, wastewater plants, and household sewage triggers eutrophication in the downstream aquatic environments ^6^. Therefore, the sustainable recovery and reuse of phosphorus is an urgent need to sustain the global food chain and other human activities, while simultaneously preserving aquatic environments.

Wastewater in particular is an abundant, widespread phosphorus sink produced by a variety of agricultural and industrial activities. Phosphorus recycling from wastewater not only would prevent further downstream ecological damage but also would lead to the development of a sustainable P bioeconomy, where the recycled phosphorus can be converted into useful, value-added P-containing materials. In addition to many well-established P removal methods, such as adsorption and chemical precipitation ^7,8^, an alternative for phosphorus recovery from wastewater was designed in the form of a biological phosphorus removal system ^9^. The biological phosphorus removal system relies on polyphosphate-accumulating organisms (PAOs) which can uptake phosphorus from wastewater and accumulate the phosphorus in the form of inorganic polyphosphate (polyP) inside cells ^10^. For example, phototrophic microalgae *Chlorella* spp. in municipal wastewater treatment plants could achieve >90% phosphorus removal ^11^. Like in *Saccharomyces cerevisiae*, the accumulated polyP in *Chlorella* spp. can reach up to 25% *Chlorella* dry weight ^12^. Additionally, the algal polyP can subsequently be extracted from cells using sonication and centrifugation in hot water for downstream application ^13^. These examples suggest that biological phosphorus removal systems can enable eco-friendly and cost-effective phosphorus removal, making them good candidates for developing the sustainable P bioeconomy.

In particular, polyP is a linear polymer of tens to thousands of phosphate residues linked by high-energy phosphoanhydride bonds, which was proposed to be a primordial energy source ^14,15^. Currently, it is known that PolyP has numerous biological functions and biomedical applications, which varies depending on the chain length (**Figure 1**); short/medium-chain polyP (10–100-mer) promotes bone regeneration ^16^, wound healing ^17,18^, and blood coagulation ^19,20^, while long-chain polyP (100–1,000-mer) are less soluble (>300-mer is insoluble)^21^ and can be used as biomolecule-carrying microdroplets that exhibit antiviral properties ^22–24^ and have molecular chaperone properties in the micromolar concentration regime ^25^. For example, the polyP 120-mer was reported to specifically bind to the angiotensin-converting enzyme 2 of human epithelial cells and thus block the SARS-CoV-2 replication ^23^. Traditionally, phosphate glass, composed of polydisperse polyP, is synthesized by heating phosphoric acid at high temperatures (>700℃) ^26^. As different applications require polyP with various specific chain lengths, the chemically synthesized polyP is then separated by length *via* liquid chromatography or fractional precipitation using organic solvents, which are resource and time-intensive processes ^27^ and also result in low yields of each polyP of specific chain length. Similar to chemical methods, the polyP purified from microalgal systems is also heterogeneous in length ^28^, which would typically require the same types of intensive separation protocols as chemically synthesized polyP if it is to be harvested and processed for practical use. Thus, for algal phosphate removal systems to be included within the sustainable P bioeconomy, the development of an environmentally friendly and non-resource-intensive method to produce polyP of a specific length, especially the valuable polyP 100-mer, is necessary.

**Figure 1.**
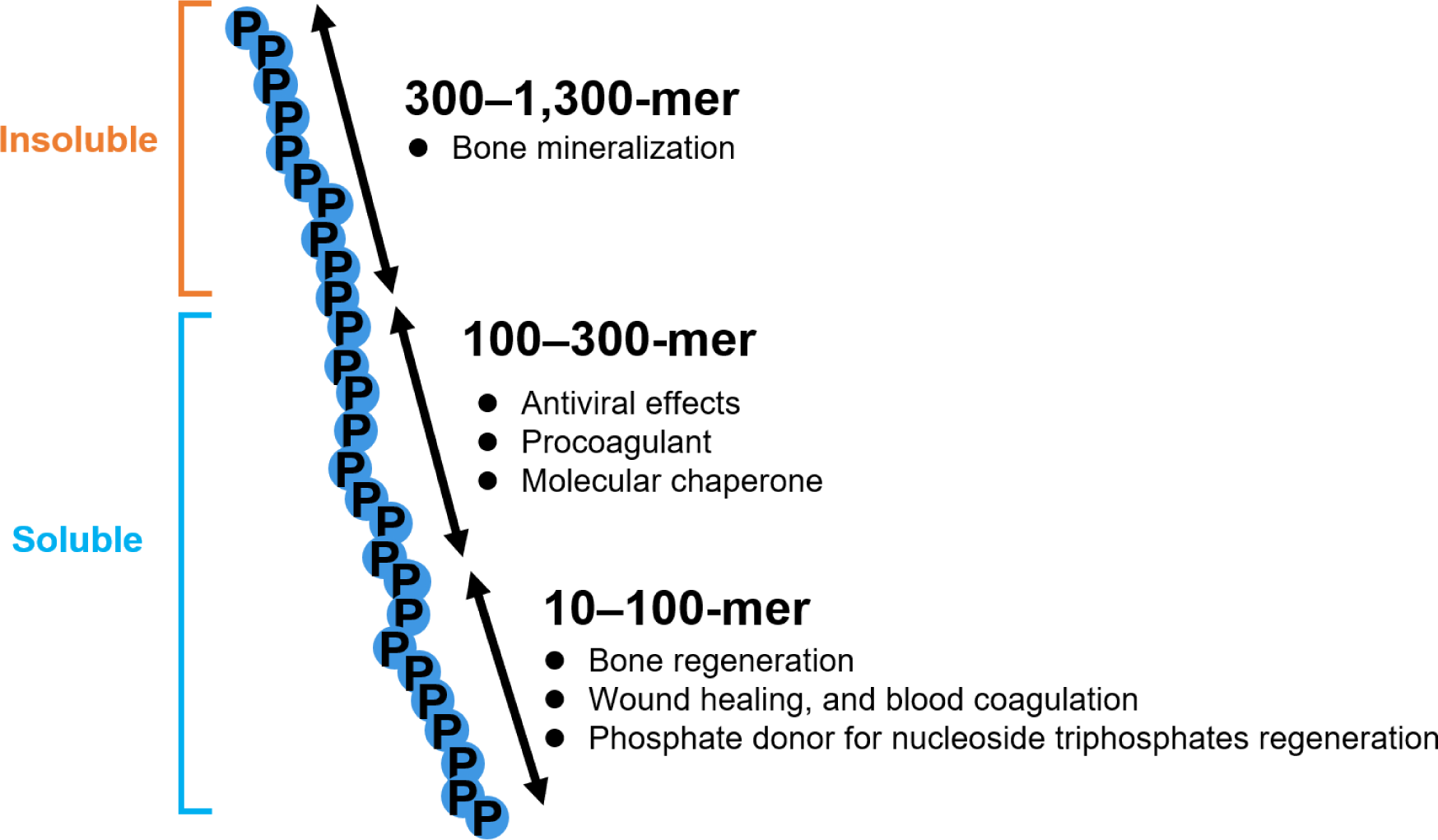
Functional diversity of polyphosphates of different lengths.

As polyP is ubiquitous in biology and because polyP function varies depending on chain length, organisms must harbor some biochemical mechanisms to produce polyP of a specific length to achieve their physiological goals. In prokaryotes, the biosynthesis and utilization of polyP are primarily mediated by polyP kinases (PPKs) with the two main families represented by PPK1s and PPK2s, which catalyze the reversible transfer of phosphate between polyP and nucleotides ^29^. Recent phylogenetic analysis has identified three subtypes of PPK2s (class I, II, and III) ^30,31^; class I and II PPK2s catalyze the polyP-driven phosphorylation of either NDP or NMP, respectively, while class III PPK2s can phosphorylate both NDP and NMP, enabling direct NTP production from NMP ^32^. Given their ability to generate NTPs from NDPs and NMPs, Class I and II PPK2s have been used for *in vitro* biosynthesis of acetone ^33^, aldehyde ^34^, thiamine phosphates ^35^, and biocatalytic regeneration of S-adenosyl-L-methionine (SAM) using polyP as phosphate donor ^36^. On the other hand, the class III PPK2s are especially useful for the cell-free protein synthesis and *in vitro* biocatalytic reactions that simultaneously require regeneration of both ATP and GTP from A(G)MP and A(G)DP. For example, the engineered highly active class III PPK2 from *Cytophaga hutchinsonii* has been applied to regenerate A(G)TP from A(G)MP *via* A(G)DP in a reconstituted cell-free protein synthesis system ^37^. In these systems, the long-chain polyP (100-mer), as opposed to the short-chain polyP at the same molar content of total orthophosphate, can significantly enhance the protein yield.

As polyP is ubiquitous in nature in all cells and because polyP function varies depending on chain length, organisms must harbor some biochemical mechanisms to produce polyP of a specific length to achieve their physiological goals; thus, we take inspiration from biology and found a biochemical mechanism to synthesize homogeneous long-chain polyP. Recently, the *Ralstonia eutropha* PPK2c was found to catalyze the direct synthesis of insoluble long-chain polyP (length undetermined) from ATP without a short-chain polyP as the primer ^38,39^. Given that *Cytophaga* PPK2 can use polydisperse polyP to phosphorylate ADP to ATP, while *Ralstonia* PPK2c can produce long-chain insoluble polyP from ATP (**Figure S1**), we aimed to harness these two PPK2 enzymes in tandem to convert polydisperse polyP in wastewater microalgae biomass into insoluble homogeneous long-chain polyP, which can be purified from the microalgal cell-lysate by one-step filtration. To prevent competition between the two phospho-transfer reactions, we used creatine as an intermediate to carry the high-energy phosphate (*i.e.*, creatine phosphate as the P-shuttle) and developed a one-pot, two-step multi-enzyme cascade for producing long-chain polyP. The polyP-rich microalgae cells were lysed to obtain cell-lysate highly enriched in polydisperse polyP (∼35 mM). After that, polyP and creatine are converted into creatine phosphate under alkaline conditions (pH 9.0) using the enzyme cascade comprising creatine kinase (CK) and polyP-consuming *Cytophaga* PPK2 (thermodynamic coupling) (**Table 1**). After adjusting the reaction mixture to neutral pH and removal of His_6_-tagged *Cytophaga* PPK2, the *Ralstonia* PPK2c was introduced to transform creatine phosphate *via* ATP to insoluble long-chain polyP (however, homogeneous instead of polydisperse) and creatine using the enzyme cascade comprising CK and polyP-synthesizing *Ralstonia* PPK2c (thermodynamic coupling). The homogeneous insoluble long-chain polyP products can then be purified by a simple one-step filtration (phase transitions), followed by non-enzymatic degradation to yield purified polyP of any length for further application.

**Table 1.** eQuilibrator-based estimation of the Gibbs free energy (Δ_r_G′^m^) of the listed enzymatic reactions under the following experimental conditions: 1 mM reactant concentration, pH 7.5, 25℃, pMg 3.0, and 0.25 M ionic strength).

| | Reaction | $\Delta_r G'^m$ (kJ/mol) |
| --- | --- | --- |
| **I** | $\text{PolyP}_{(n)} + \text{ATP} \rightarrow \text{PolyP}_{(n+1)} + \text{ADP}$ | ~0 |
| **II** | $\text{ADP} + \text{Creatine phosphate} \rightarrow \text{ATP} + \text{Creatine}$ | -12.2 |
| **I + II** | $\text{PolyP}_{(n)} + \text{Creatine phosphate} \rightarrow \text{PolyP}_{(n+1)} + \text{Creatine}$ | -12.2 |

## Results

To develop the sustainable P bioeconomy process, we used discharge samples from a local piggery wastewater treatment system as a substrate for microalgae cultivation and polyP production (**Figure 2A**). This wastewater was first sterilized by heat and then the microalgae *Chlorella vulgaris* was cultivated under nitrogen-deficient conditions to induce the assimilation of phosphorus in the form of polyP ^40^. After cultivation, toluidine blue O (TBO) was used to live-stain the microalgal cells and visualize the intracellular polyP *in vivo*. Optical microscopy observations showed the accumulation of small purple-stained particles, approximately 1 µm in diameter, which were likely highly enriched in polyP (**Figure 2B**). The polyP-accumulating microalgal biomass was then collected by centrifugation and lysed by sonication, followed by heating at 100°C (**Figure 2C**). This resulted in microalgal cell-lysates containing up to 35 mM polyP (**Figure 2D**) and demonstrating that the algal polyP can be produced using simple cultivation and extraction processes. The cell-lysate polyP exhibited solid particle-like structures that are heterogeneous in size, which can be directly observed through epifluorescence microscopy after DAPI (4,6-diamidino-2-phenylindole) staining (**Figure 2E**). Moreover, the polyP in the cell-lysate appeared to be polydisperse in length based on the results of TBE-Urea polyacrylamide gel electrophoresis analysis (**Figure 2F**). We hypothesized that the polydisperse polyP in algal cell-lysates can be reduced and elongated in length using enzymatic activities of the *Cytophaga* class III PPK2 and *Ralstonia* PPK2c, respectively, to produce insoluble long-chain polyP (**Figure S1**).

**Figure 2.**
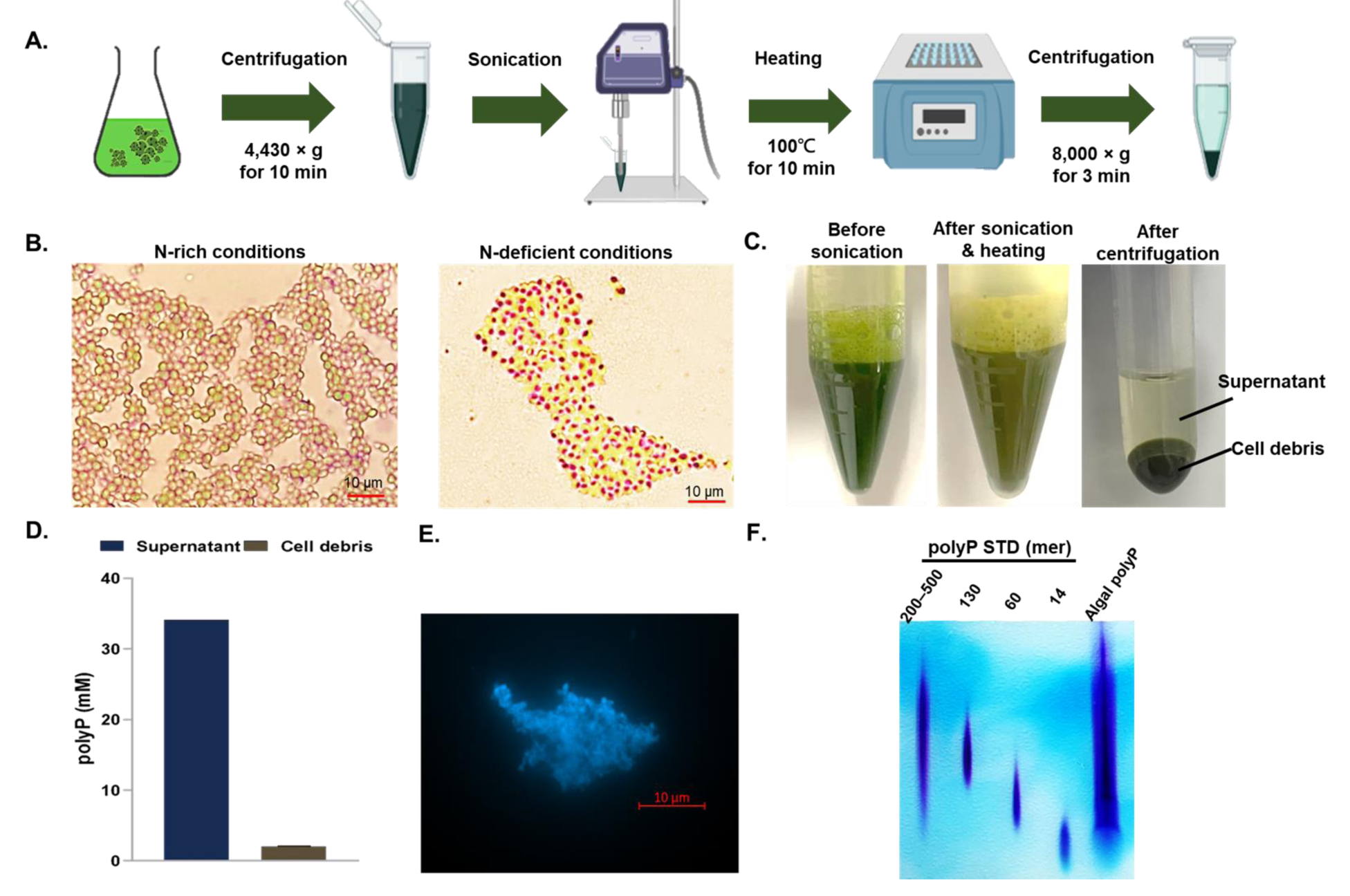
Microalgae cultivation and partial fractionation of the accumulated polyphosphate (polyP). **(A)** The overall scheme for producing polydisperse algal polyP. **(B)** PolyP accumulation in *Chlorella vulgaris* cultivated in sterilized wastewater under nitrogen-deficient conditions. The intracellular polyP was visualized *in vivo* by TBO staining and analyzed by optical microscopy. **(C)** Production of the polyP-rich cell-lysate (supernatant) from microalgal biomass *via* sonication, heating, and centrifugation. **(D)** The soluble polyP concentrations in the supernatant and the cell debris (measured by the TBO assay). Error bars represent the standard deviation from three experimental replicates. **(E-F)** DAPI-stained epifluorescent microscopy analysis (**E**) and TBE-Urea polyacrylamide gel electrophoresis (6%, w/v) analysis (**F**) of the granular polydisperse polyP aggregates.

The next step for the proposed sustainable P bioeconomy includes converting the polydisperse polyP in microalgal cell-lysates to another P-containing molecule (creatine phosphate) for the downstream synthesis of homogeneous long-chain polyP; however, the prerequisite of this step is that the polydisperse polyP in the microalgal cell-lysate can serve as the substrate of *Cytophaga* PPK2, similar to what is possible with commercial polyP 25-mers (**Figure S1**), so that the high-energy phosphate can be completely transferred to the downstream P-carrier. The theoretical product of each *Cytophaga* PPK2-mediated phospho-transfer reaction is ATP and polyP with one less unit in the chain (polyP_(n)_ + ADP → polyP_(n-1)_ + ATP). To measure the reaction kinetics for stoichiometric analysis, we coupled the *Cytophaga* PPK2-mediated ATP production process to an NADP reduction process driven by an enzyme cascade consisting of hexokinase (HK) and glucose-6-phosphate dehydrogenase (G6PD) (**Figure 3A**). In the coupled HK/G6PD enzyme cascade, glucose is first converted into glucose-6-phosphate by HK using one ATP, which is then converted into dehydro-glucose-6-phosphate, along with the reduction of one NADP to produce one NADPH, which can be observed through λ_340 nm 37_. Upon incorporation of the HK/G6PD cascade to the *Cytophaga* PPK2-mediated ATP production process, we observed NADPH accumulation over time upon progression of this reaction in the microalgal cell-lysate (**Figure 3B**); stoichiometric analysis also confirmed that the NADPH production (*i.e.*, ATP regenerated) is equivalent to polyP consumption, suggesting that all high-energy phosphate contained within polyP was transferred fully to ATP (**Figure 3C**).

**Figure 3.**
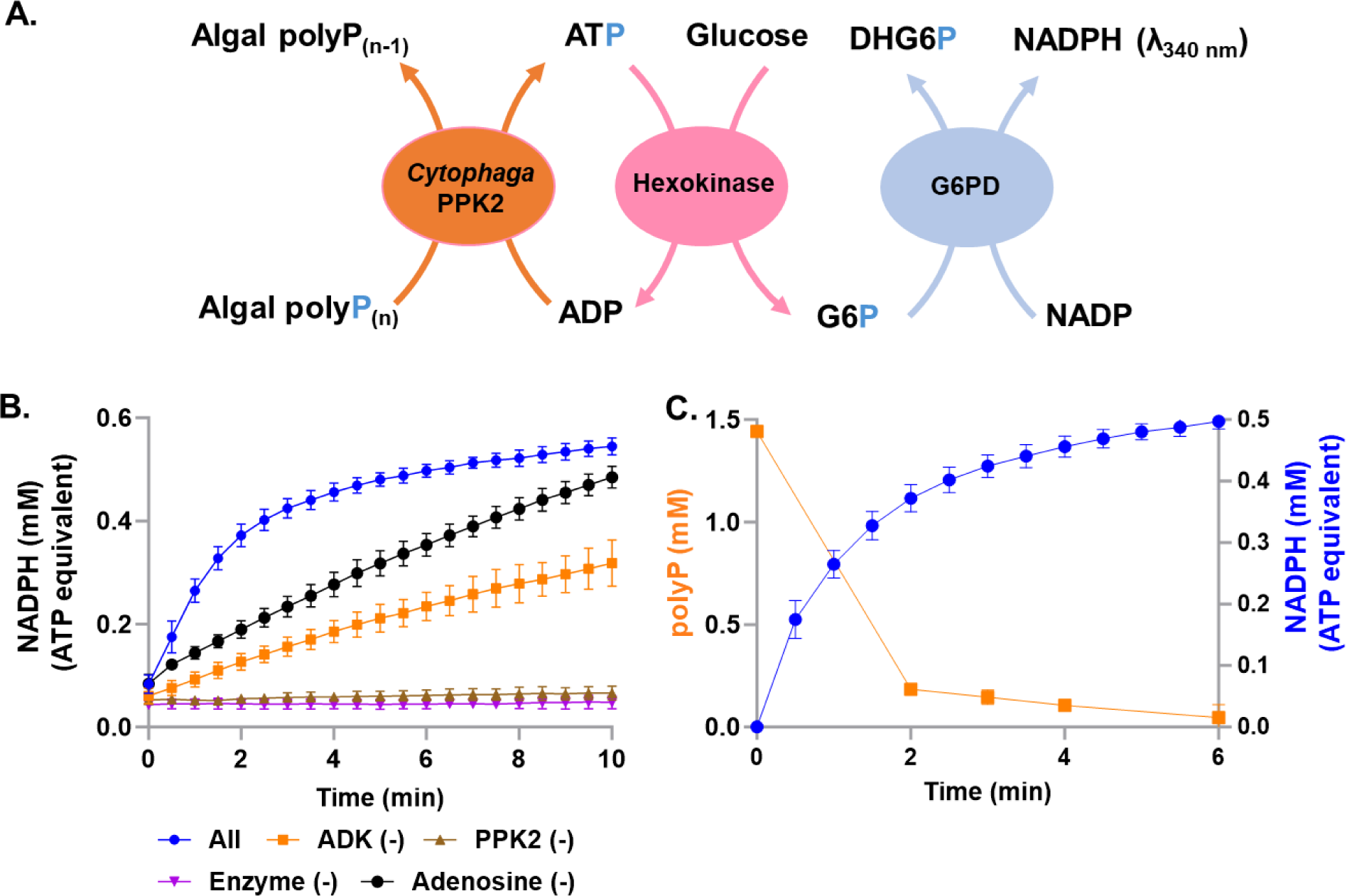
*Cytophaga* PPK2-based ATP regeneration using polydisperse polyP in microalgal cell-lysate. **(A)** Schematic diagram showing the enzymatic cascade of the *Cytophaga* class III PPK2 and HK-G6PD-coupled NADPH production assay. HK; hexokinase, G6PD; glucose-6-phosphate dehydrogenase, DHG6P; dehydroglucose-6-phosphate. **(B)** PolyP-based ATP regeneration monitored by ATP-dependent NADH production (λ_340 nm_) using G6PD-HK. **(C)** Stoichiometric analysis of *Cytophaga* PPK2-dependent polyP consumption and HK-G6PD-coupled NADPH production. The concentrations of the consumed polyP and produced NADPH was monitored through the TBO assay and at λ_340 nm_, respectively. The error bars represent the range and the data points represent the average from two independent experimental replicates.

We then chose creatine phosphate as the P-carrier for downstream synthesis of insoluble long-chain polyP (**Figure 4A**; **Table 1)**, as eQuilibrator-based free energy calculations suggest that CK-mediated phospho-transfer from ATP to creatine is thermodynamically favorable at basic pH (**Figures 4B** and **S2A**) ^41^. Given the previous demonstration that P from algal polyP can be fully converted to ATP, complete phospho-transfer from the polydisperse polyP to creatine *via* ATP in the microalgal cell-lysate is plausible. On the other hand, the CK-mediated phospho-transfer from creatine phosphate to ADP (the reverse reaction) is thermodynamically favorable at neutral pH (**Figure S2B**). Therefore, by modulating the pH of the microalgal cell-lysate, one can first convert the polydisperse polyP and creatine into creatine phosphate *via* ATP (polyP_(n)_ + creatine → polyP_(n-1)_ + creatine phosphate) using polyP-consuming *Cytophaga* PPK2 and CK at basic pH, and later can convert creatine phosphate back into long-chain polyP (but insoluble) and creatine using CK and polyP-synthesizing *Ralstonia* PPK2c *via* ATP at neutral pH (polyP_(n)_ + creatine phosphate → polyP_(n+1)_ + creatine).

**Figure 4.**
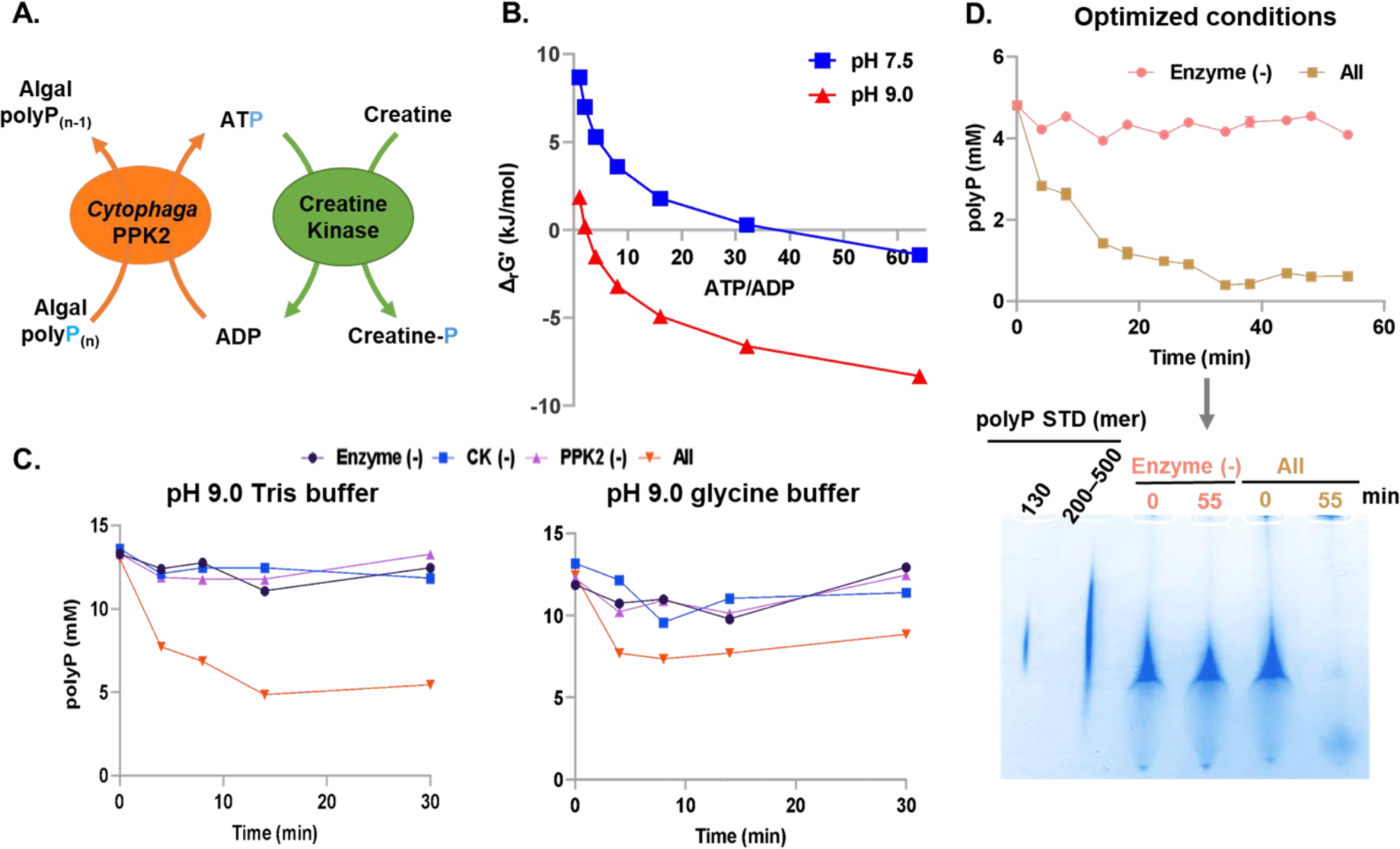
Conversion of polydisperse algal polyP into creatine phosphate *via* ATP by the enzymatic cascade comprising CK and *Cytophaga* PPK2. **(A)** Schematic diagram showing the PPK2-CK enzyme cascade. **(B)** eQuilibrator-based thermodynamic calculations of creatine phosphorylation at circumneutral (pH 7.5) or alkaline (pH 9.0) pH. **(C)** Time-dependent creatine phosphate production by the PPK2-CK cascade in Tris or glycine buffer at pH 9.0. The production of creatine phosphate was monitored by the consumption of the polyP *via* TBO assay. **(D)** Time-dependent creatine phosphate production by the PPK2-CK cascade under optimized conditions (Tris (pH 9.0), Mg^2+^ (10 mM), creatine (50 mM), algal polyP (5 mM)). The reactions were conducted with and without *Cytophaga* PPK2. The nearly complete consumption of polyP was verified *via* quantitative TBO measurements (top) from TBE-Urea polyacrylamide gel electrophoresis analysis (bottom).

Thus, we sought to validate the proposed phospho-transferase cascade for creatine phosphate production from the polydisperse algal polyP **(Figure 4A)**. Using free energy calculations as a guide (**Figure S3**), we first optimized the conditions of this two-enzyme PPK2/CK cascade. Based on our experimental analysis, 10 mM Mg^2+^ at pH 9.0 in Tris buffer resulted in the greatest polyP concentration decrease (*i.e.*, creatine phosphate production) (**Figures 4C** and **S3AB**). We next observed that 50 mM creatine concentration was optimal for both polyP consumption and creatine phosphate conversion (to prevent data misinterpretation solely based on the polyP consumption, HPLC analysis was used to measure creatine phosphate conversion in parallel) (**Figures S3CD**). Finally, 5 mM algal polyP also resulted in the greatest amount of creatine phosphate conversion (∼4.75 mM; 95% yield) (**Figure S3E**). Using these optimized conditions (10 mM Mg^2+^, 50 mM creatine, and 5 mM algal polyP at pH 9.0 in Tris buffer), nearly complete polyP consumption was observed (**Figure 4D**), and thus the subsequent creatine phosphate-producing reactions were all performed using these conditions.

Next, with creatine phosphate generated from algal polyP, we sought conditions to transfer the high-energy phosphate on creatine phosphate to build a growing polyP chain. Thus, we then applied a two-enzyme cascade containing CK and *Ralstonia* PPK2c in HEPES-K at neutral pH (7.0); the high-energy phosphate from creatine phosphate will be transferred to ADP *via* CK (regenerating ATP), while the *Ralstonia* PPK2c would transfer the high-energy phosphate on the regenerated ATP onto a growing chain of polyP, producing long-chain polyP (**Figure 5A**). In this enzyme cascade, ATP serves as a P shuttle, transferring the high-energy phosphate from creatine phosphate to the growing polyP chain. We first sought to optimize the yield of the CK/PPK2c cascade at different ATP concentrations **(Figure 5B)**. Although it may seem that a higher initial ATP concentration may lead to more production of polyP, eQuilibrator-based free energy calculations revealed that increasing ATP concentration would also inhibit the phospho-transfer from creatine-phosphate to ADP (**Figure S2B**) and thus would block the elongation of polyP chain. Experimentally, the data suggested that 3.5 mM ATP can maximize long-chain polyP production (**Figures 5C and S4**). In the absence of creatine phosphate (but with added ATP), nearly no long-chain polyP was produced, suggesting the importance of creatine phosphate to drive the aforementioned exergonic phospho-transfer (thermodynamic coupling). Therefore, we successfully converted the polydisperse polyP into long-chain polyP *via* creatine phosphate in the microalgal cell-lysate.

**Figure 5.**
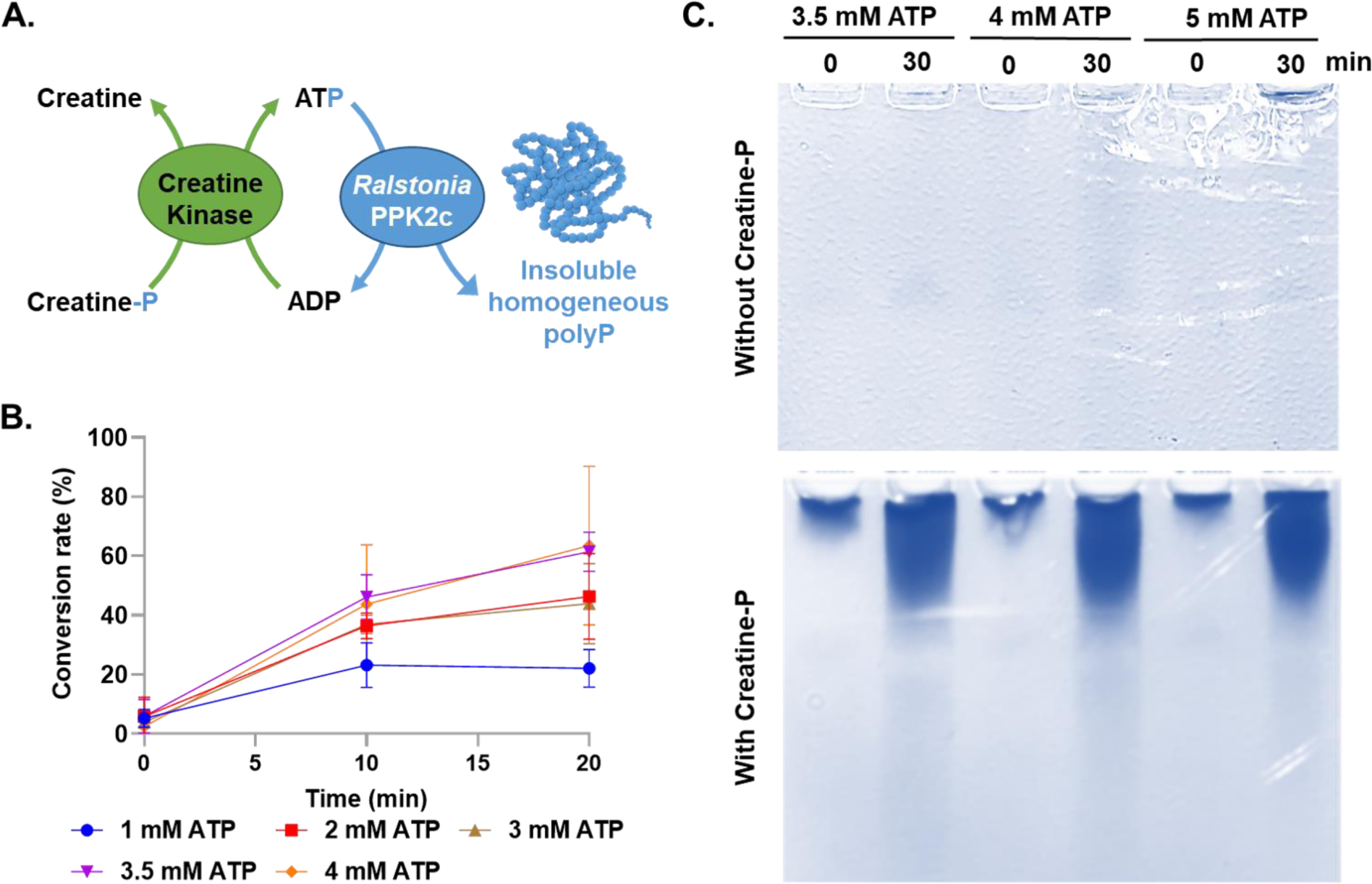
Conversion of creatine phosphate into homogeneous insoluble long-chain polyP *via* ATP by the enzymatic cascade comprising CK and *Ralstonia* PPK2c. **(A)** Schematic diagram showing the two-enzyme cascade comprising CK and *Ralstonia* PPK2c for homogeneous insoluble long-chain polyP production. **(B)** Time-dependent long-chain polyP production by the CK-PPK2c cascade in HEPES-KOH buffer (pH 7.5) with varying ATP concentrations. **(C)** TBE-Urea polyacrylamide gel electrophoresis analysis of homogeneous long-chain polyP production. The reaction was conducted with and without creatine phosphate, along with variations in ATP concentration. Error bars represent the standard deviation and the data points represent the mean from three independent experimental replicates.

Next, given that both enzymatic cascades (*Cytophaga* PPK2-CK and CK-*Ralstonia* PPK2c) were shown separately to be effective, we then sought to perform the entire reaction in a one-pot, two-step fashion for greater throughput and scalability. Specifically, we first performed the creatine phosphate-producing cascade (*Cytophaga* PPK2-CK) at pH 9.0 (**Figure 6A**). After that, we added Ni Sepharose to remove the *Cytophaga* PPK2 and adjusted the reaction pH to neutral (optimal for transformation of creatine phosphate to ATP by CK) (**Figure 6B**). Finally, *Ralstonia* PPK2c was added to transform the produced creatine phosphate to ATP (via CK) and then to the long-chain polyP (using PPK2c) (**Figure 6C**). However, our experimental analysis revealed that the *Cytophaga* PPK2-CK cascade and the CK-*Ralstonia* PPK2c require different buffer systems; specifically, none of the buffer systems tested (Tris, carbonate-bicarbonate buffer, and glycine buffer) at the required pH range (pH 7.0–9.0) allow both cascade reactions to occur. Thus, we reasoned that rather than using either buffer in purified form, a mixture of both buffers at an intermediate pH may facilitate both cascades, albeit possibly with sub-optimal efficacy for both cascades. Among all conditions tested, a HEPES-K:Tris ratio of 8:1 resulted in the greatest long-chain polyP production (**Figure S5A**). Further optimization of the one-pot, two-step reaction in 8:1 HEPES:Tris revealed that the concentrations of pH, ATP, CK, and creatine phosphate of pH 7.0, 3.5 mM, 0.1 mg/mL, and 5 mM, respectively, resulted in the greatest long-chain polyP yield (90%) (**Figures S5B–E**). In parallel, we observed a nearly complete conversion of the creatine phosphate into long-chain polyP and creatine by the CK-*Ralstonia* PPK2c cascade under the same assay conditions but in the HEPES-K buffer (**Table 2**), suggesting that the mixed buffer is indeed sub-optimal for the CK-*Ralstonia* PPK2c cascade. However, considering that the *Cytophaga* PPK2-CK cascade requires completely different conditions, the final conditions, while perhaps sub-optimal for either or both cascades, can still produce long-chain polyP at high yield (90%), sufficient for demonstration of the efficacy one-pot two-step process.

**Figure 6.**
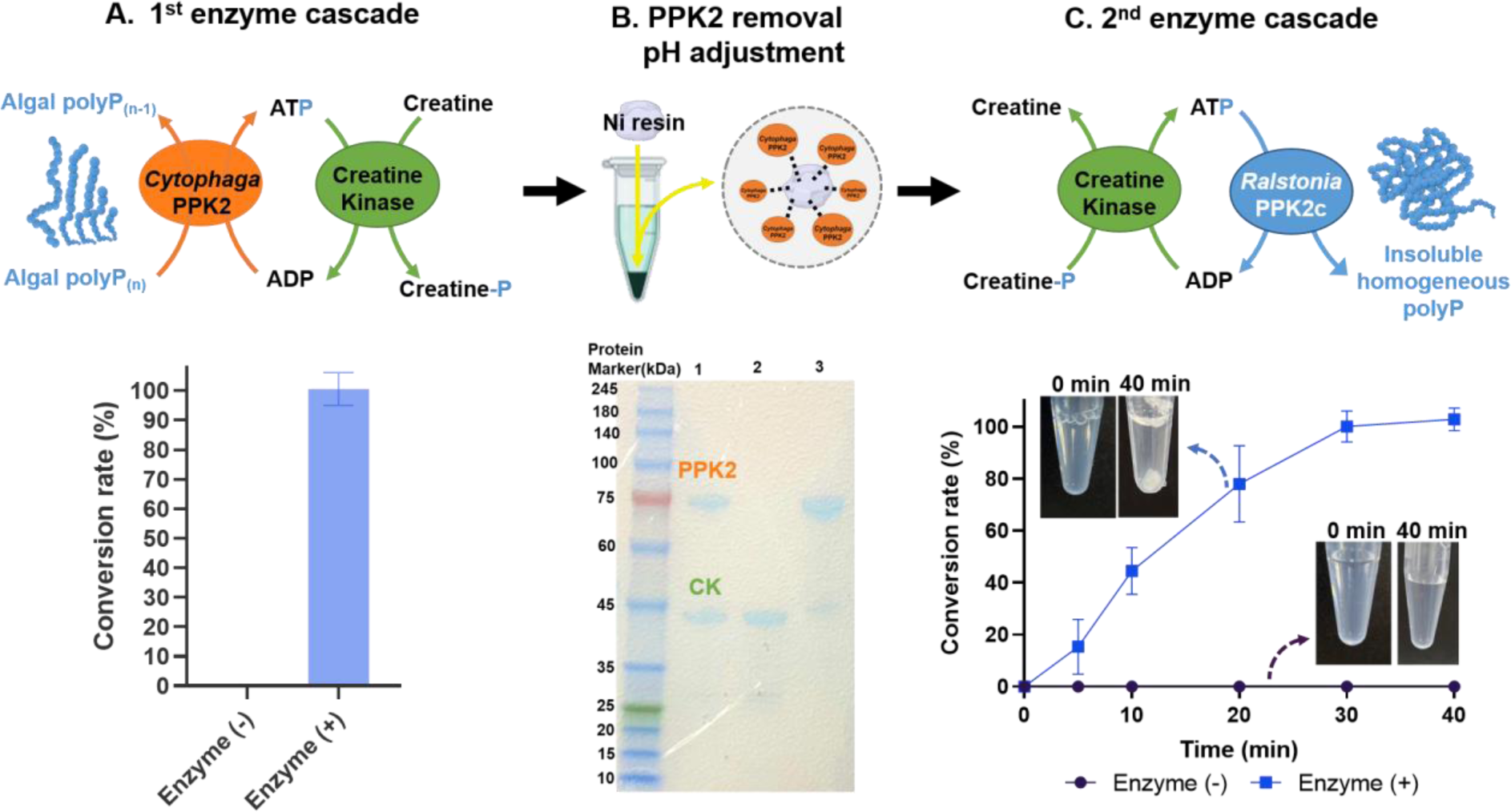
One-pot, two-step enzymatic synthesis of homogeneous insoluble long-chain polyP from polydisperse algal polyP. **(A)** Conversion of polydisperse algal polyP into creatine phosphate *via* the *Cytophaga* PPK2-CK cascade. **(B)** The removal of His-tagged *Cytophaga* PPK2 from the microalgal cell-lysate (verified by SDS-PAGE) using the Ni-chelating resin. 1: the cell-lysate with both *Cytophaga* PPK2 and CK; 2: the cell-lysate after *Cytophaga* PPK2 removal by a Ni-chelating resin; 3: the elution of the Ni-chelating resin used for *Cytophaga* PPK2 removal. A trace amount of CK was also co-eluted. **(C)** Conversion of creatine phosphate into homogeneous insoluble long-chain polyP solids *via* the CK-*Ralstonia* PPK2c cascade. The conversion rates of the insoluble long-chain polyP synthesis reaction with the mixed buffer system were calculated at different time points. The reactions were conducted with and without *Ralstonia* PPK2c. Error bars represent the standard deviation and the data points represent the mean from three independent experimental replicates.

**Table 2.** The output summary of each step of the one-pot, two-step polyphosphate (polyP) synthesis.

| Step | Substrates | Products | Yield (%) | Residues and Byproducts |
| --- | --- | --- | --- | --- |
| **A. Synthesis of creatine phosphate** | Algal polyP,<br>Creatine | Creatine phosphate | 95% | Creatine,<br>Algal polyP (< 5-mer) |
| **B. Enzyme removal/pH adjustment** | N.A. | N.A. | 99% | N.A. |
| **C. Synthesis of long-chain polyP** | Creatine phosphate,<br>ATP | Insoluble long-chain polyP 1,300-mer | 90% | Creatine |
| **D. Filtration** | N.A. | N.A. | 99% | N.A. |
| **E. Hydrolysis of long-chain polyP** | Long-chain polyP 1,300-mer | polyP 100–1,300-mer | 90% | P<sub>i</sub> |

We also observed insoluble material produced after the one-pot, two-step enzymatic cascade (**Figure 6C**), which we conjectured was the long-chain polyP product itself. If this assumption is correct, then the insoluble long-chain polyP products would be quite high in molecular weight (polyP > 300-mer is generally insoluble), and thus we first filtered the products using a 100-kDa (∼1,000-mer) cutoff centrifugal filter and then measured the polyP concentrations of both the flow-through and the remainder on the filter. The polyP products appeared to be all “ultra” long-chain polyP because nearly no polyP was detected in the flow-through (**Figure 7A**; **Table 2**). This is in contrast to the polydisperse polyP in microalgal cell-lysate before the enzymatic catalysis, which has roughly equal concentrations of polyP of sizes larger and smaller than 100 kDa (**Figure 2F**). TBE-Urea polyacrylamide gel electrophoresis analysis and HPLC analysis both suggested that the polyP products were highly homogeneous and in the 1,300-mer unit range (**Figure S6A–B**). Previous studies have focused on long-chain polyP (700-mer; some studies refer to these as “super long-chain” polyP); however, our synthesized homogeneous product, which could be purified from the microalgal cell-lysate *via* a one-step filtration, is nearly twice as long compared to the longest commercially available polyP.

**Figure 7.**
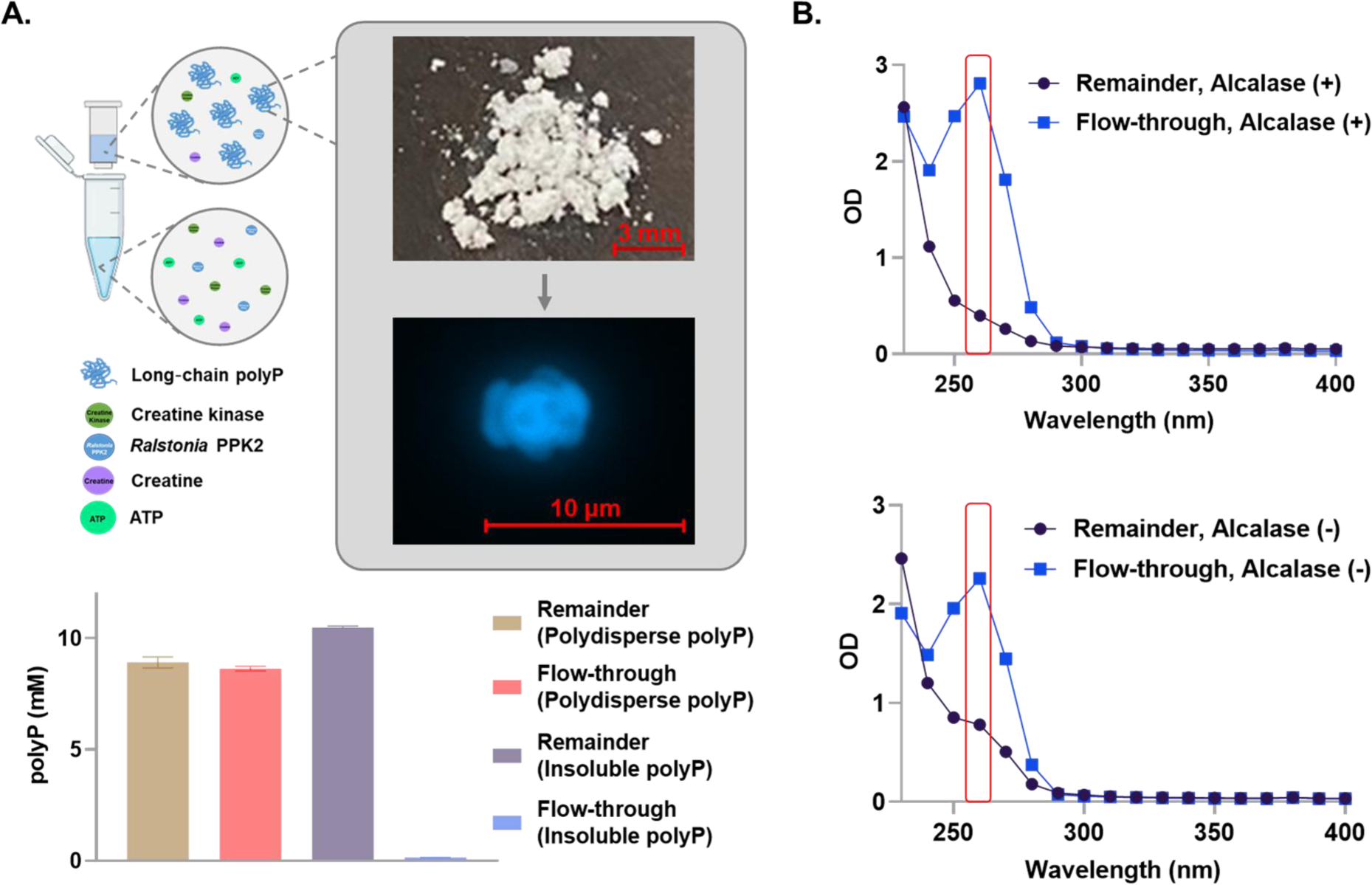
Purification of long-chain polyP using a membrane filter after the protease digestion. **(A)** The solutions containing the polydisperse algal polyP or the insoluble homogeneous long-chain polyP obtained from the one-pot, two-step enzymatic cascades were subjected to filtration through a 100-kDa filter. PolyP concentrations in the remainder and flow-through fractions were quantified by the TBO assay. **(B)** Removal of small molecules (ATP, creatine, and salts) and proteins from the remainder fraction (verified by UV-Vis analysis). The reaction mixture containing insoluble long-chain polyP was subjected to filtration before and after the proteolysis treatment.

Although homogeneous long-chain polyP has been produced *via* our one-pot, two-step enzymatic cascades, the product could potentially contain some byproducts or contaminants, such as nucleic acids and peptides, that would inhibit downstream use or processing for industrial purposes. We thus further subjected the microalgal cell-lysate containing the polyP 1,300-mer product to a protease treatment and filtration by a 0.45-µm filter for polyP purification. Consistently, ATP and proteins (indicated by λ_260-280 nm_) were nearly completely washed away in the remainder based on UV-Vis and SDS-PAGE analysis (**Figures 7B and S6C; Table 2**), suggesting effective purification of the polyP 1,300-mer product. After filtration, we then dried the remainder, which resulted in a white powder that fluoresced after DAPI-staining, confirming its composition to be of polyP (**Figure 7A**).

While the goal of this study was to convert polydisperse polyP in wastewater microalgae biomass into insoluble and homogeneous long-chain polyP, we next wondered whether the developed process could lead to other value-added products aside from the polyP 1,300-mer. As mentioned previously, polyPs of different lengths have very different functional properties, and the ability to acquire polyPs of different lengths is of particular value. Prior to this study, industrial production methods for polyP of different chain lengths required extensive chromatographic processes or gradient organic solvent precipitation for fractionation from polydisperse polyP (produced from alkaline-treated phosphate glass), both of which are time-, resource-, cost-, and organic waste-intensive. Thus, we finally resolved to explore the possibility of producing shorter homogeneous polyP from the polyP 1,300-mer. We first decided to subject the polyP 1,300-mer to enzymatic treatment by exopolyphosphatase (PPX) ^42^. However, it appeared that rather than decreasing the length of the polyP product over time, the polyP concentration instead decreased over time, with very little change in its chain length (**Figure S7A**). Moreover, *Cytophaga* PPK2 treatment of polyP 1,300-mer also resulted in a similar result (**Figure S7B**). We attribute this to the fact that PPX and *Cytophaga* PPK2 likely degrade single polyP chains fully before moving on to the next chain. Such an enzymatic degradation strategy was not amenable to our goals.

We thus decided to search for a non-enzymatic aqueous strategy that did not degrade single polyP chains fully. We subjected the polyP 1,300-mer (5 mM) to non-enzymatic digestion at neutral pH (7.5) and different temperatures with Mg^2+^ (5 mM) as the catalyst for the non-enzymatic polyP digestion, along with 5 mM ethylenediaminetetraacetic acid (EDTA) (Mg^2+^ chelator) to minimize non-enzymatic polyP endo-cleavages. Our data revealed that the length of the polyP products was slightly reduced in a time-dependent manner at 70°C (**Figure 8A**); however, even after 4 hours of incubation, the size of the polyP products was still quite large (and the chain length remained much higher than the polyP 200–500-mer marker). Thus, we decided to increase the reaction temperature to 95°C. Over just one hour, the length of the polyP was reduced in a time-dependent manner efficiently, ultimately reaching a length on the order of 100-mer (while passing through the entire range of polymer lengths between 100–1,300-mer) (**Figure 8B**). Moreover, the overall polyP concentration remained greater than 90% after the non-enzymatic hydrolytic process, confirming this process to be efficient with minimal loss of polyP product (**Figure 8C**, **Table 2**). Thus, large quantities of purified polyP of specific lengths between 100–1,300-mer can be produced from the polyP 1,300-mer obtained from our enzymatic method, something not possible with any other synthetic polyP method developed to date. As mentioned previously, our prior study revealed that PPK2 is more efficient in utilizing a polyP 100-mer than commercial short-chain polyP (25–65-mer) for ATP regeneration (at the same phosphate molar content). Thus, to demonstrate the added value of the non-enzymatic hydrolytic polyP 100-mer product while confirming its activity, we used the polyP 100-mer product to perform the *Cytophaga* PPK2-based ATP regeneration process. Indeed, we observed more efficient ATP regeneration in the assays using the polyP 100-mer than those using the commercial short-chain polyP (**Figure 8D**), suggesting the added value of the ultimately produced polyP 100-mer. We also note that other than the 100-mer, polyP of other lengths that are non-enzymatically generated from the homogeneous 1,300-mer, especially those between 100-mer and 300-mer, could also be used for applications such as those in biomedicine (**Figure 1**). We believe that we have demonstrated that integration of the entire chemo-enzymatic system presented herein has resulted in a sustainable P bioeconomy platform valorizing low-value biomass waste to produce high-value products for various applications.

**Figure 8.**
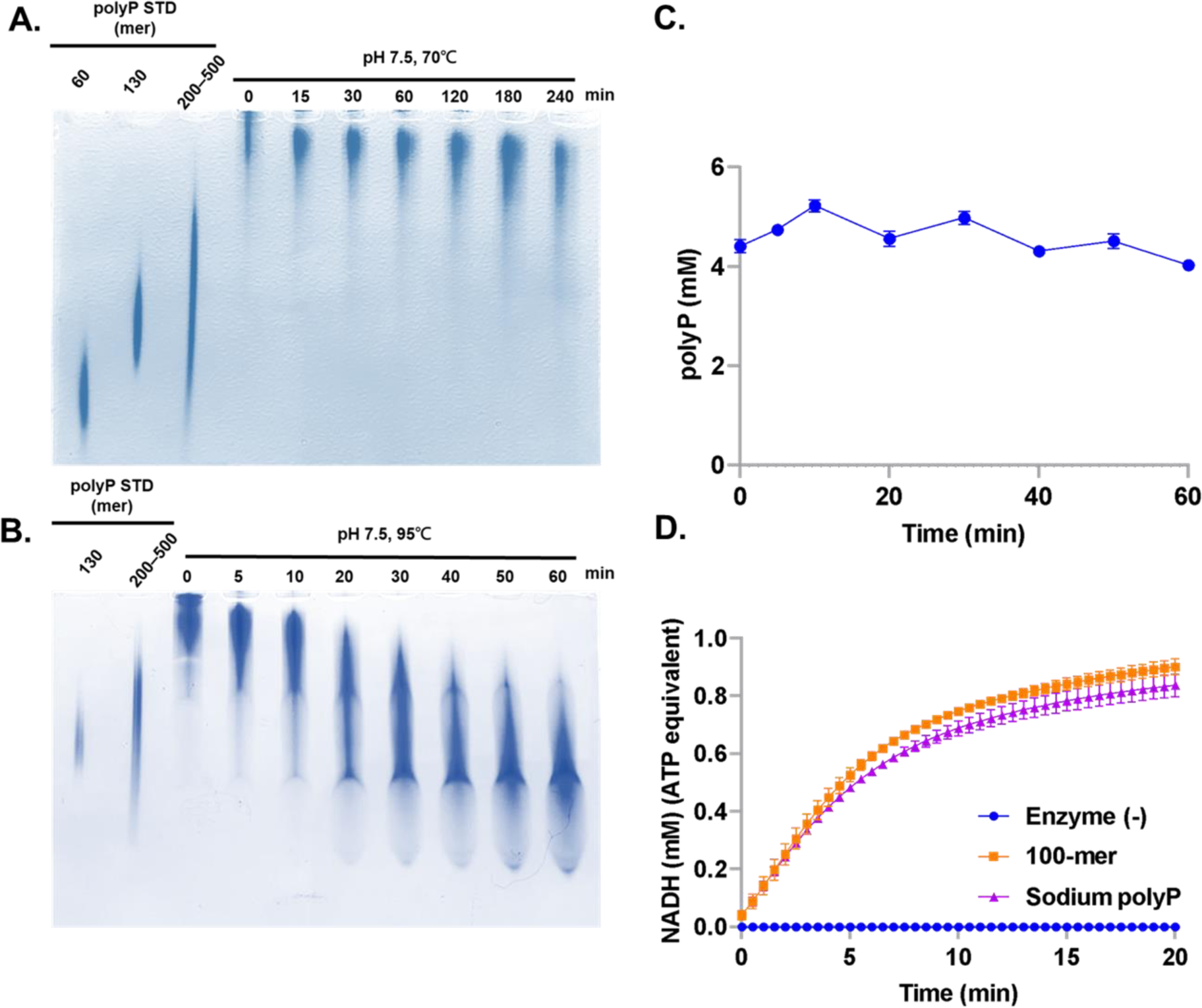
Time-dependent thermo-digestion of a homogeneous polyP 1,300-mer by non-enzymatic hydrolysis. **(A-B)** The polyP 1,300-mer was incubated at **(A)** 70℃ and **(B)** 95℃ and at pH 7.5, along with 5 mM Mg^2+^ and 5 mM ethylenediaminetetraacetic acid. The reaction mixtures collected at different time points were analyzed by TBE-Urea polyacrylamide gel electrophoresis, along with commercial polyP standards as a reference for the lengths. **(C)** The total concentration of polyP (based on the molar content of orthophosphate) during the time-dependent thermo-digestion was monitored by TBO assay. **(D)** HK-G6PD-mediated NADH production, which was coupled to *Cytophaga* PPK2-mediated ATP regeneration; commercial short-chain polyP and purifiedpolyP 100-mer product was the high-energy phosphate donor (normalized to the same molar content of orthophosphate). Error bars represent the standard deviation and the data points represent the mean from three independent experimental replicates.

## Discussion

In this study, we devised an efficient enzyme cascade to sustainably produce polyP 1,300-mer from wastewater microalgal biomass (or from commercial short-chain polyP). This technology simultaneously purifies wastewater to avoid eutrophication of downstream aquatic environments (**SDG 6**), while also mitigating the global phosphorus deficit and producing high-value biomedical materials (in the form of shorter polyP of specific length between 100-mer and 1,300-mer) following non-enzymatic hydrolysis (**SDG 3**). From a biochemical standpoint, the success of this technology results from the unusual properties of (***i***) CK that allow a pH-based modulation of the direction of polyP-ATP phospho-transfer and (***ii***) *Cytophaga PPK2* and *Ralstonia* PPk2c that allow a two-step back-and-forth polyP phospho-transfer. However, this technique also succeeds due to a unique phase-transition property of the polyP reactants and products. In biology, phase transitions have often been employed to circumvent thermodynamic limitations, which can direct and inhibit the reversibility of bio-polymerization reactions to accumulate high concentrations of polymerization products in cells ^14^, as is also observed in the case of polyP accumulation in the *Chlorella* cells (**Figure 2B**). We thus used the same principles to drive the enzymatic synthesis of solid long-chain polyP from fully soluble shorter-chain polyP, where the phase-transition of the polyP products from soluble to insoluble leads to the favorability of the forward polyP synthesis process in solution. Moreover, the solidity of the long-chain polyP 1,300-mer products facilitates a streamlined, one-step polyP purification procedure *via* simple filtration to yield purified polyP that could be applied to downstream use.

The presented microalgal cultivation and extraction procedures at the lab scale also have the potential to be up-scaled to industrial levels through collaboration with industry. While microalgal biomass collection, sonication-based cell disruption, and heating seem to be easily scalable, the centrifugation step required for the insoluble microalgal polyP separation from other cell debris could be one hurdle in the development of the initial steps of any large-scale procedure in the future due to capacity limitations in the total volume of microalgae that can be centrifuged. Therefore, future development of techniques that can facilitate both protease/lipase-based cell lysis to allow us to access the microalgal polyP and membrane-based filtration to separate the microalgal polyP from other cell debris at a large scale would be required to bring the long-chain polyP synthesis method into development at a larger-scale. Similarly, the bio-enzymatic procedures to convert polydisperse algal polyP into insoluble polyP 1,300-mer *via* creatine phosphate have currently been designed as a one-pot, two-step cascade at the lab scale. Future optimization that allows the enzymatic conversion process to increase in efficiency and sustainability, as well as procedures to upscale this process, would be essential to facilitate long-chain polyP at the commercial scale. For example, the use of magnetic nanoparticles to immobilize the His-tagged enzymes could bypass the need for centrifugation at a large scale and allow the re-use of the enzymes. Moreover, further investigations into a “panacean” buffer system that could accommodate the required catalytic conditions for all enzymatic members would streamline the procedure to allow all required phospho-transfers without any loss in yield in one-pot, which could be up-scaled more readily than the current design. Finally, upon acquisition of the polyP 1300-mer products, we developed a time-dependent non-enzymatic digestion method to produce purified polyP of any length shorter than 1,300-mer over 1 hour as a more effective alternative to the currently used commercial procedure, which involves low-yield fractionation of a polydisperse polyP mixture to acquire polyP of different lengths and is both cost- and time-consuming. The unexpected stability of the polyP 100-mer as compared to longer polyPs in the presence of Mg^2+^ and EDTA at 95℃ and circumneutral pH allows for high-yield polyP 100-mer production (∼90%) from the polyP 1,300-mer.

Given that the *Chlorella* spp. where the microalgal insoluble polyP starting material are acquired is regarded as **G**enerally **R**ecognized **a**s **S**afe (GRAS) by the USA Federal Drug Administration (FDA), we believe that the value-added polyP products of various lengths reliably produced by our novel procedure are suitable to be used in biomedicine. In particular, polyP products of specific lengths can be used in bone stitches (300–1300-mer), as antivirals (100–300-mer), or as drug delivery vessels (10–100-mer). In particular, the future discovery of the unexplored biological functions or medical applications of purified polyP products of lengths greater than 700-mer (other than bone materials) could also result in greater value for our system. Nevertheless, we also demonstrated that the polyP 100-mer can act as the energy stock to drive ATP-dependent cell-free protein synthesis and biocatalysis, and thus we believe that any polyP product, including those greater than 700-mer, has the potential to be applied in the same way. Furthermore, the intermediate creatine phosphate synthesized using the algal polyP could also be used as medicine for heart failure, cardiac surgery, and skeletal muscle hypertrophy ^43,44^.

Altogether, the catalytic processes established in this study facilitate a sustainable P bioeconomy platform that can valorize microalgal biomass to produce value-added polyP products at the lab scale. However, we believe that a large-scale global sustainable P bioeconomy is crucial to solving the imminent loss of all global phosphate sources in the next 70 years. Thus, we expect that upon scale-up and further development, the scale of the sustainable P bioeconomy platform will increase to allow the production of large amounts of high-value polyP materials that are essential for biotechnology and medicine. The development of such a system allowing sustainable large-scale production of such polyP materials from highly abundant microalgal biomass is essential for decreasing humankind’s reliance on a diminishing global resource. In particular, as microalgae are abundant not only in wastewater, but any bodies of water, an initial application of our polyP synthesis technique in global regions with coasts or rivers that undertake significant phosphorus mineral mining activities (an environmentally harming activity) would help those regions to divest from economic reliance on phosphorus mineral mining (**SDG 9**). The subsequent establishment of a sustainable P bioeconomy in other regions lacking phosphorus minerals would help to drive the establishment of local, self-sustainable polyP material production, thereby reducing impacts both of phosphate mineral mining as well as environmental and financial costs related to constant shipping and acquisition of polyP materials.

## Acknowledgments

The authors acknowledge Toshikazu Shiba and other employees of RegeneTiss for providing EX-polyP as standard samples, as well as performing HPLC gel-filtration analysis of the synthesized polyP 1,300-mer. P.-H.W. is supported by the National Science and Technology Council of Taiwan (112-2628-E008-005). T.Z.J. is supported by the Japan Society for the Promotion of Science (JSPS) (Grant-in-aid 21K14746) and the Mizuho Foundation for the Promotion of Science. K.F. is supported by an ELSI research grant and NINS Astrobiology Center grant (AB301003 and AB311001).

## Author contributions

T.Z.J. and P.-H.W. conceptualized the project and designed experiments. Y.-H.L., S.N., F.-I.,Y., and P.-H.W. performed experiments. All authors contributed to data analysis and interpretation. Y.-H.L., S.N., T.Z.J., and P.-H.W. wrote the manuscript with support from all authors.

## Declaration of interests

The authors declare no competing interests.

## Data and code availability

The published article and the supplementary data include all datasets generated or analyzed during this study.

## Experimental procedures

Methods for **free energy calculation**, **protein expression and purification**, **electrophoretic analysis of polyP using TBO-stained TBE-Urea polyacrylamide gel electrophoresis**, **High-Performance Liquid Chromatography (HPLC) analysis**, enzymatic digestion and elongation of polyP using *Cytophaga* PPK2 and *Ralstonia* PPK2c, and **determination of the length distribution of the generated polyP 1,300-mer** are described in the **Supplemental Information**. The detailed information and kinetic properties of the enzymes used in this study are available in **Supplemental Tables S1** and **S2**. The **SDS-PAGE analysis of the enzymes used in this study**, **HPLC chromatogram of creatine and creatine phosphate**, **the optical standard curves of polyP and NADH**, and **the geographic information of the phosphate-rich wastewater sampling site** are available in the **Appendix**. Chemicals and reagents are purchased from Sigma-Aldrich (St. Louis, MO, USA) unless specified otherwise.

### Quantification of polyP using the toluidine blue O (TBO) method

PolyP was quantified by a metachromatic assay with the TBO method using commercial polyP (sodium polyP) as a standard. The TBO method is based on the concentration-dependent decrease in λ_630 nm_ by the metachromatic reaction of TBO with polyP ^45^. Briefly, sample solution (5 µL) was mixed with TBO assay solution (250 µL; 15 µg/mL) and acetic acid (0.1 N) at room temperature ^46^. Then, λ_630 nm_ was measured for the TBO-treated sample in a microplate spectrophotometer for 10 min (Molecular Devices/Spectra Max® iD3, San Jose, CA, USA). The λ_630 nm_ was later converted into polyP concentration based on standard curves derived from the different concentrations of commercial sodium polyP standards. The standard curves of polyP concentrations are available in the **Appendix**.

### Microalgae cultivation under nitrogen-deficient conditions

Microalgae *Chlorella vulgaris* (*C. vulgaris*) was purchased from the Bioresource Collection and Research Center (Hsinchu, Taiwan) and was cultivated in heat-sterilized wastewater collected from the discharge of a local piggery wastewater treatment plant with continuous daylight exposure (**Appendix**). *C. vulgaris* was cultivated in 2-L Erlenmeyer flasks containing the sterilized wastewater (1 L; pH adjusted to neutral) at room temperature with continuous shaking (200 rpm) for aeration and to prevent microalgae from sticking to the bottom of the flask as previously described ^47^.

### Epifluorescence microscopic detection of polyP

PolyP was detected by epifluorescence microscopy as previously described ^38^. Briefly, polyP granules were stained with DAPI (4′,6-diamidino-2-phenylindole) (0.1 mg/mL in distilled H_2_O) for at least 10 min and the stained granules were visualized by epifluorescence microscopy with oil at a 1,000 x magnification (ZEISS/AXIOSKOP 2, Oberkochen, Germany).

### In vivo polyP visualization using TBO staining

*C. vulgaris* cells were air-dried and heat-fixed on a glass slide (76 × 26 mm; Thickness 1.2–1.5 mm). Intracellular polyP granules were then stained with TBO (15 mg/L) for 10 min by submerging the whole glass slide (containing the fixed cells) into TBO solution. The slide was then gently washed with double distilled H_2_O, followed by air drying for 15 min and subsequent observation by an optical microscope at a 100 x magnification (Olympus CX21FS1, Shinjuku, Tokyo, Japan).

### C. vulgaris cell lysis and partial polyP purification

The *C. vulgaris* cells were disrupted and partially purified as previously described ^45^. *C. vulgaris* biomass was collected by centrifugation at 4,430 × g for 10 min at room temperature and then resuspended in buffer (HEPES-K (pH 7.0; 20 mM), KCl (0.15 M), and ethylenediaminetetraacetic acid (5 mM)) at a pellet to buffer ratio of 1:3. The cells were lysed *via* ultrasonication for 20 min (3 s on and 3 s off) and the cell-lysate containing polyP was subsequently incubated at 100℃ for 10 min, followed by centrifugation at 8,000 × g for 3 min at room temperature to separate the cell debris from the supernatant containing the polydisperse polyP. The polyP concentration within the supernatant and the initial microalgal wastewater were quantified by the TBO method (see above). The supernatant containing polyP was stored at –80°C for further use in subsequent experiments.

### ATP regeneration using heterologous algal polyP

Polydisperse polyP in the microalgal cell-lysate was converted into ATP *via* the ATP regeneration cascade. The reaction mixtures (200 µL) contained Tris (pH 7.0; 100 mM), Mg^2+^ (10 mM), polyP (1.5–10 mM), adenosine (1–3 mM), *Cytophaga* PPK2 (0.08 mg/mL). The reaction was initiated by the addition of PPK2. ATP production was monitored at 37℃ for 10 min by both (***i***) the time-dependent consumption of polyP using the TBO method (see above) and (***ii***) the hexokinase/glucose-6-phosphate dehydrogenase (Roche, Basel, Switzerland)-coupled NAD^+^ reduction process (λ_340 nm_) as described previously ^37^.

### Enzymatic synthesis of creatine phosphate from polydisperse polyP in microalgal cell-lysate

A two-enzyme cascade comprising *Cytophaga* PPK2 and rabbit creatine kinase (CK) (Sigma-Aldrich) was applied to sequentially convert the algal polyP into creatine phosphate *via* ATP. The optimized reaction mixtures (200 μL) contained Tris (pH 9.0; 0.1 M), MgSO_4_ (10 mM), polyP (10 mM), creatine (50 mM), ATP (1 mM), N-acetyl-L-cysteine (2 mM), *Cytophaga* PPK2 (0.3 mg/mL), and CK (0.03 mg/mL); different conditions, including pH 8.0, 5 mM and 15 mM MgSO_4_, and 10–40 mM creatine were also tested, but the reported reaction conditions are the optimized conditions used for all subsequent experiments. The reaction was initiated by the addition of *Cytophaga* PPK2 and CK, and the formation of creatine phosphate was monitored at 30℃ for 30 min by the consumption of the algal polyP using the TBO method (see above) as well as HPLC analysis.

### Enzymatic synthesis of homogeneous polyP 1,300-mer

Another two-enzyme cascade comprising *Ralstonia* PPK2c (polyP-synthesizing) and rabbit CK was used to sequentially convert creatine phosphate into homogeneous polyP 1,300-mer *via* ATP. The formation of the polyP 1,300-mer was monitored by the TBO method (see above). The reaction mixtures (200 μL) contained (HEPES-K (pH 7.0; 90 mM), Tris (pH 7.0; 10 mM), MgSO_4_ (10 mM), creatine phosphate (5 mM), ATP (3.5 mM), PPK2c (0.5 mg/mL), and CK (0.1 mg/mL); different ATP concentrations from 1–5 mM were also tested, but the reported reaction conditions are the optimized conditions used for all subsequent experiments. The reaction was initiated by the addition of CK and *Ralstonia* PPK2c at 30℃, and the formation of the polyP 1,300-mer was monitored *via* the time-dependent decrease in λ_630 nm_ using the TBO method.

### Degradation of homogeneous polyP 1,300-mer by non-enzymatic hydrolysis

The synthesized polyP 1,300-mer in the microalgal cell-lysate was collected by filtration using a 0.45-µm MF-Millipore® membrane filter paper (Burlington, Massachusetts, USA) along with a vacuum pump. The remainder was washed by ddH_2_O until the intensity of λ_265 nm_ (indicative of N(M/D/T)P) and λ_280 nm_ (indicative of protein/polypeptide) of the flowthrough decreased to background levels. After resuspension of the reaction by adding 300 µL HEPES-K buffer (25 mM, pH 7.5), the reaction mixture (MgSO_4_ (5 mM), EDTA (5 mM), and polyP 1,300-mer (5 mM)) was subjected to time-dependent hydrolysis at 95°C.

## Notes

### Competing Interest Statement

The authors have declared no competing interest.

