## Supplemental Materials for "One-pot chemo-enzymatic synthesis and one-step recovery of homogeneous long-chain polyphosphates from microalgal biomass"

### Supplemental Experimental Procedures

##### *Free energy calculation*

The Gibbs free energy for each biochemical reaction was computed utilizing the eQuilibrator tool (https://equilibrator.weizmann.ac.il/) [1]. The Gibbs free energies of the reactions at physiological standards ($\Delta rG'^{m}$) were approximated with the following parameters: reactant concentrations of 1 mM, ionic strength of 0.25 M, 1.0 mM $Mg^{2+}$, pH at 7.5, and a temperature of 25°C. The eQuilibrator API was used to estimate reaction Gibbs energies (ΔG′) at an ionic strength of 0.1 M and 30°C, employing varying conditions as delineated below. For the creatine phosphorylation reaction, initial estimation of ΔG′ was carried out across different pH values (7.0–10.0) and ATP concentrations (0.1–5.0 mM) under the specified experimental conditions: creatine at 15 mM, ADP at 0.15 mM, creatine phosphate at 15 mM, and Mg2+ at 10 mM. This was aimed at identifying the optimal pH and ATP concentration. Subsequently, the impact of diverse $Mg^{2+}$ concentrations (0.1–100 mM) on ΔG′ for the creatine phosphorylation reaction was evaluated under varying pH conditions (4.0–10), while retaining ATP at 10 mM. Furthermore, the influence of varying creatine concentrations (0.1–50 mM) on the ΔG′ of the creatine phosphorylation reaction was assessed at pH 9.0, using differing creatine phosphate concentrations (5.0/10 mM), and maintaining the following experimental conditions: $Mg^{2+}$ at 10 mM, ATP at 0.7 mM, and ADP at 0.3 mM. For the reverse reaction (creatine de-phosphorylation), ΔG′ was determined at pH 7.0 along with varying ATP concentrations (0.1–10 mM), under the specified experimental conditions: $Mg^{2+}$ at 10 mM, ADP at 0.3 mM, creatine at 5 mM, and creatine phosphate at 0.8 mM.

***Protein Expression and Purification***

For standard expression and purification of recombinant enzymes used in this study (*Cytophaga* PPK2, *Ralstonia* PPK2c, and *E. coli* exopolyphosphatase (**Table S1**)), recombinant *E.coli* BL21(DE3) cultures were cultivated under aerobic conditions at 37°C in a shaker containing Terrific Broth at 200 rpm until a specific optical density was reached ($OD_{600\ nm}$ 0.6), followed by induction with isopropyl β-D-1-thiogalactopyranoside (0.4 mM), and then by overnight incubation at 16°C. Recombinant proteins were purified to at least 95% homogeneity using immobilized metal affinity chromatography (Ni Sepharose High-Performance resin) as previously described [2], and protein purity was analyzed by SDS-PAGE (**Appendix**). In brief, cells were collected by centrifugation at 8,000 × g for 10 min at 4°C and resuspended in binding buffer (HEPES (4-(2-hydroxyethyl)-L-piperazineethanesulfonic acid)-Na buffer (pH 7.5; 25 mM), NaCl (0.3 M), glycerol (5%, w/w), and imidazole (5 mM)). The cells were then homogenized *via* sonication in an ice bath for 10 min (1 s on and 2 s off). The lysate was centrifuged at 20,000 × g for 20 min at 4°C and the supernatant was passed through a glass gravity column containing 0.5 mL Ni Sepharose™ High-Performance resin (GE Healthcare, Chicago, IL, USA). After washing with wash buffer (HEPES-Na buffer (pH 7.5; 25 mM), NaCl (0.3 M), glycerol (5%, w/w), and imidazole (25 mM)), proteins were eluted with 0.4 mL of elution buffer (HEPES-Na buffer (pH 7.5; 25 mM), NaCl (0.3 M), glycerol (5%, w/w), and imidazole (300 mM)). The protein stock solutions were flash-frozen in liquid nitrogen, stored at −80°C, and thawed immediately before use. The N-$His_6$ tag was retained for all experiments. All subsequent enzyme assays were conducted in duplicate or triplicate as indicated in each respective figure.

##### *Electrophoretic analysis of polyP using TBO-stained TBE-Urea polyacrylamide gel electrophoresis*

Electrophoretic analysis of polyP was performed following an established protocol described previously [3]. The reaction mixtures containing polyP were pre-mixed with 2x sample buffer (2x Tris-Borate-EDTA (TBE), urea (7 M), and bromophenol blue (0.1 g/L)) and loaded onto a separating gel consisting of urea (7 M), acrylamide (6%, v/v), and 1x Tris-Borate-EDTA (TBE). Electrophoresis was performed in a 1x TBE buffer bath for 25 min at 200 V at room temperature (25°C). The gel was stained in TBO solution containing urea (0.05%, w/v), methanol (25%, v/v), and glycerol (5%, w/w) for 10 min, and was de-stained for 1 h in an identical solution but without TBO. After de-staining, the magenta-colored bands on the TBE-Urea gel represent TBO-stained polyP. The size range of the polyP was estimated by comparison to polyP standards (chain lengths of 14, 60, 130, and 200–500 (average chain length: 243); provided by Toshikazu Shiba of RegeneTiss).

##### *High-Performance Liquid Chromatography (HPLC) analysis*

Quantification of creatine phosphate and adenosine triphosphate by HPLC was performed following a protocol described previously with some modifications [4]. Briefly, the separation of creatine phosphate and adenosine triphosphate in the reaction mixture was performed on a 150 × 4.6 mm Luna® C18(2) 100 Å, LC column packed with 5-µm particles (Phenomenex, USA) while simultaneously recording $\lambda_{210\,nm}$, indicative of creatine phosphate. The acquired $\lambda_{210\,nm}$ values were later converted to creatine phosphate concentration based on a standard curve derived from the different concentrations of the corresponding chemicals. The mobile phase consisted of buffer A ($Na_2HPO_4$ (20 mM)) and buffer B ($Na_2HPO_4$ (200 mM), and acetonitrile (10%, v/v)), all adjusted to pH 5.0 with phosphoric acid (2 M) and filtered through a 0.22-μm membrane filter. The total regeneration time for the HPLC column is 20 min. Aliquots of 10 μL were injected into the instrument by autosampler, and the subsequent elution was performed at a flow rate of 0.4 mL/min in buffer A for 2 min, followed by a 1:1 gradient elution with buffers A and B up to 8.5 min, and additionally with buffer B up to 10 min. The column was cleaned by washing with 80% methanol after each run for 10 min, and was re-equilibrated with buffer A for 1 min.

***Enzymatic digestion/elongation of polyP using Cytophaga PPK2/Ralstonia PPK2c*** *Cytophaga* PPK2 and *Ralstonia* PPK2c were used to degrade and elongate commercial polyP (~25-mer), respectively. The degradation of polyP was conducted within reaction mixtures (200 μL) containing HEPES-K (pH 7.5; 100 mM), $Mg^{2+}$ (5 mM), polyP (~25-mer; 1 mM), ADP (8 mM), and *Cytophaga* PPK2 (0.1 mg/mL), while polyP elongation (200 μL) was conducted in reaction mixtures with ATP (8 mM) and *Ralstonia* PPK2c (0.5 mg/mL). The reactions were initiated by the addition of *Cytophaga* PPK2/*Ralstonia* PPK2c at 30℃. The length of the resulting polyP was monitored by TBE-Urea polyacrylamide gel electrophoresis.

***Determination of the length distribution of the generated polyP 1,300-mer***

Toshikazu Shiba of RegeneTiss (Kunitachi, Tokyo, Japan) performed gel filtration chromatography *via* HPLC, using EX-polyP (available from RegeneTiss) of various sizes (14-mer, 60-mer, and 130-mer) as a standard, as according to experimental details described in previous publications [5]. We similarly applied polyacrylamide gel electrophoresis using the same standards to confirm the length of the homogeneous long-chain polyP products in parallel.

### Supplemental Figures.

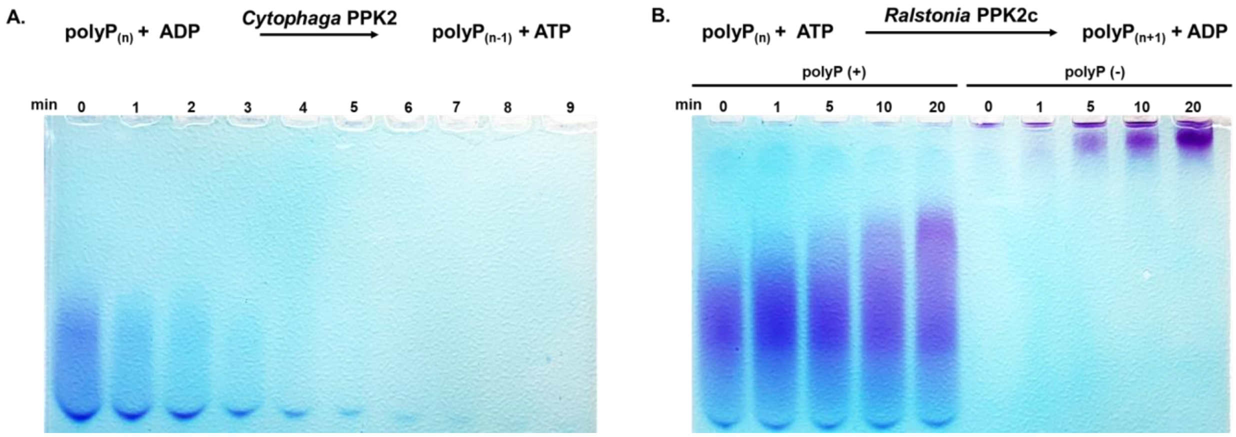


**Figure S1. Electrophoretic analysis of time-dependent polyP 25-mer mixture degradation or elongation using *Cytophaga* PPK2 or *Ralstonia* PPK2c, respectively.** Related to Figs. 3–5. **(A)** *Cytophaga* PPK2 and **(B)** *Ralstonia* PPK2c were used for the degradation or elongation of the commercial polyP 25-mer. The reaction mixtures from different time points were analyzed by TBE-Urea polyacrylamide gel electrophoresis.

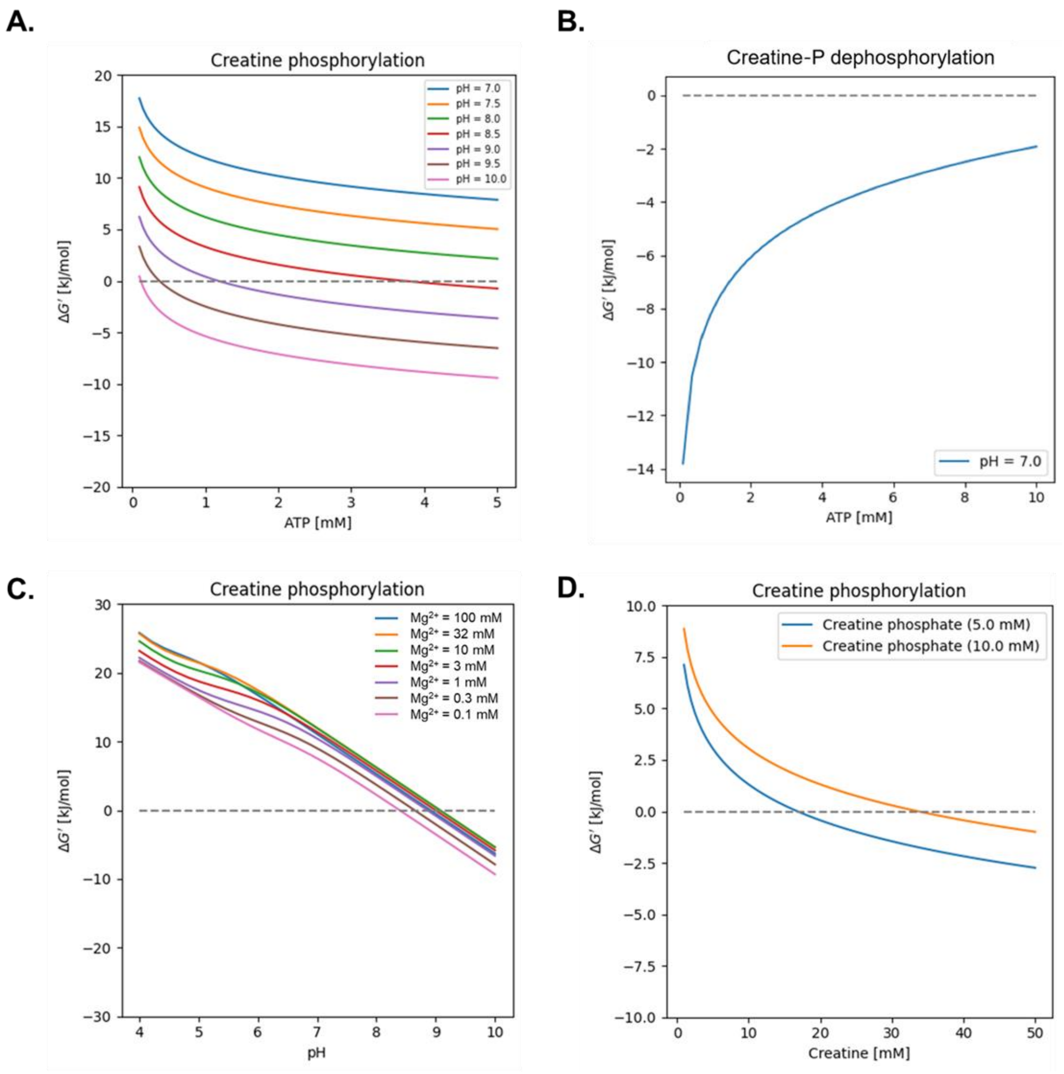


**Figure S2. eQuilibrator-based estimation of the reaction gibbs free energy (ΔG′) of the enzymatic reactions.** Related to Figs. 4 and 5. **(A)** The ΔG′ of the creatine phosphorylation reaction was estimated at different pH and concentrations of ATP under the following experimental conditions: ionic strength (0.1 M), creatine (15 mM), ADP (0.15 mM), creatine phosphate (15 mM), $Mg^{2+}$ (10 mM), and 30℃. **(B)** The ΔG′ of the creatine phosphate dephosphorylation reaction was estimated at pH 7.0 along with different concentrations of ATP under the following experimental conditions: ionic strength (0.1 M), $Mg^{2+}$ (10 mM), ADP (0.3 mM), creatine (5 mM), creatine phosphate (0.8 mM), and 30℃. **(C)** The ΔG′ of the creatine phosphorylation reaction was estimated at different pH and concentrations of $Mg^{2+}$ under the following experimental conditions: ionic strength (0.1 M), creatine (15 mM), ADP (0.15 mM), and creatine phosphate (15 mM), ATP (10 mM), and 30℃. **(D)** The ΔG′ of the creatine phosphorylation reaction was estimated at pH 9.0 at different concentrations of creatine phosphate along with different concentrations of creatine under the following experimental conditions: ionic strength (0.1 M), $Mg^{2+}$ (10 mM), ATP (0.7 mM), ADP (0.3 mM), and 30℃.

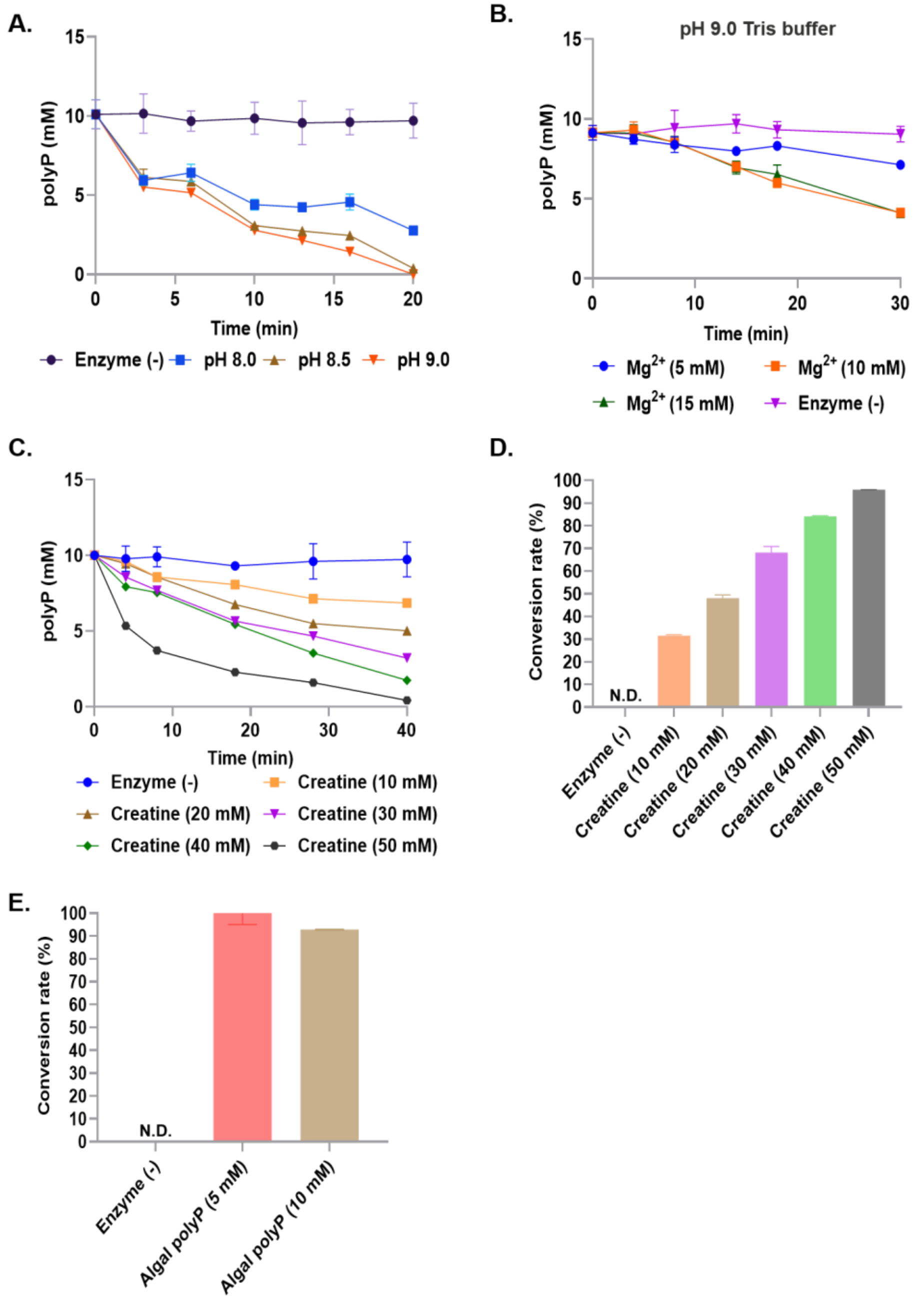


**Figure S3. Optimization of the CK-PPK2c cascade reaction for creatine phosphate production from the algal polyP and creatine.** Related to Fig. 4. Creatine phosphate production by the PPK2-CK cascade was evaluated based on polyP consumption. The PPK2-CK cascade reactions were conducted at different pH (**A**), different concentrations of $Mg^{2+}$ (**B**), and different concentrations of creatine **(C)** in Tris (pH 9.0). The conversion rates of polyP to creatine phosphate in the reaction mixture with different concentrations of creatine (**D**) and the algal polyP (**E**) were determined using HPLC. Error bars represent means ± SEM from independent experiments in triplicate.

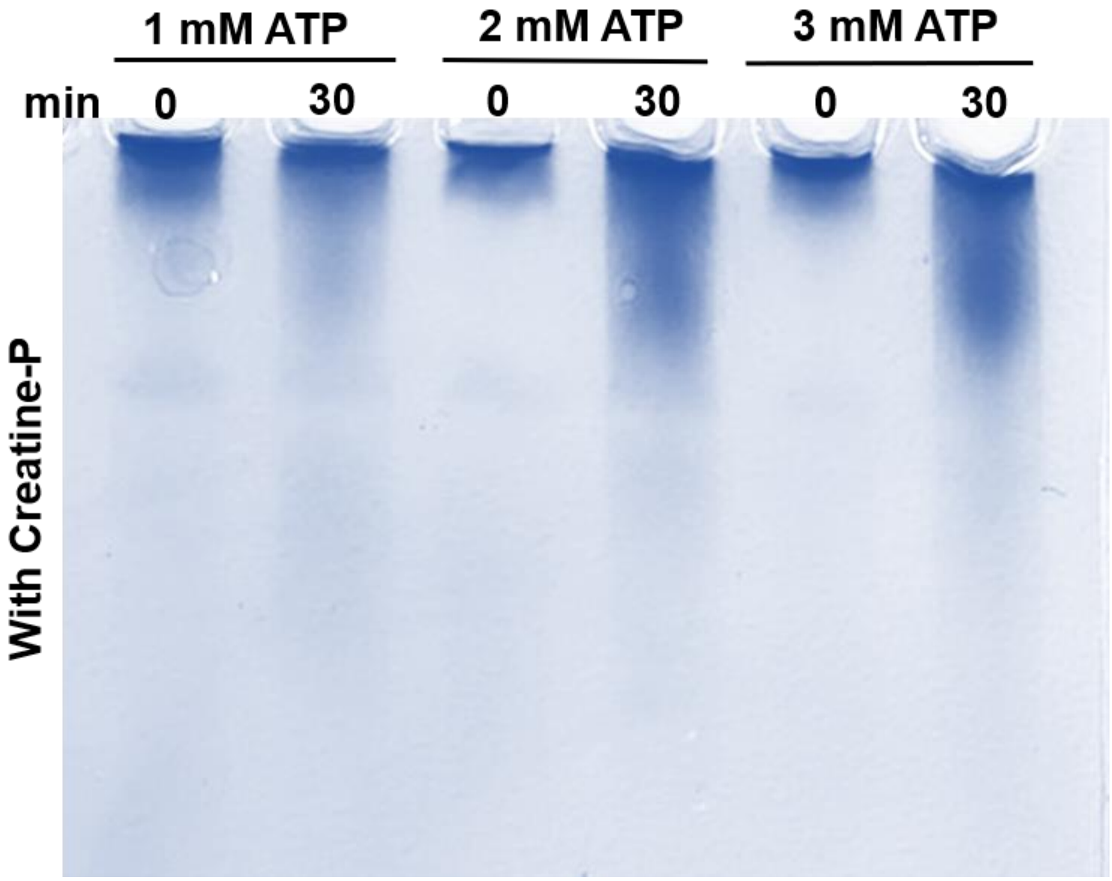


**Figure S4. Supplemental figure of Figure 5C**. Related to Fig. 5. TBE-Urea polyacrylamide gel electrophoresis analysis of homogeneous long-chain polyP production at different ATP concentrations with creatine phosphate.

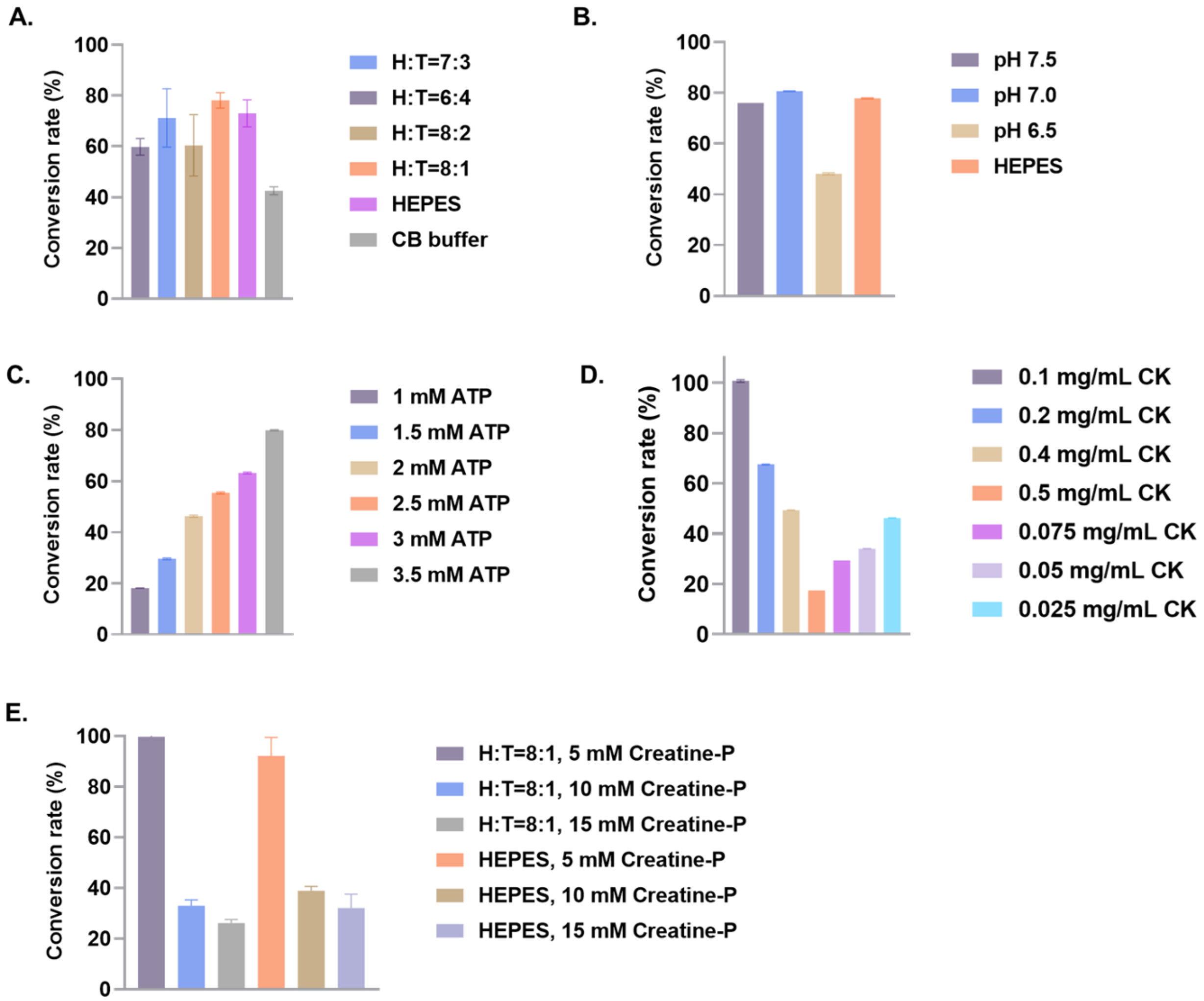


**Figure S5. Optimization of the CK-PPK2c cascade reaction for the one-pot, two-step enzymatic synthesis of insoluble long-chain polyP.** Related to Figs. 5 and 6. The CK-PPK2c cascade reactions were conducted in the mixed buffers with different ratios of HEPES and Tris (**A**). The same cascade reaction was then conducted in 8:1 HEPES:Tris at different pH (along with reactions in HEPES at pH 7.0) (**B**) and at different concentrations of ATP (**C**), CK (**D**), and creatine phosphate (along with reactions in HEPES at the same creatine phosphate concentrations) (**E**). The conversion rates of creatine phosphate to polyP in the CK-PPK2c cascade reaction under different conditions were determined using high-performance liquid chromatography. Values represent means ± SEM from independent experiments in triplicate.

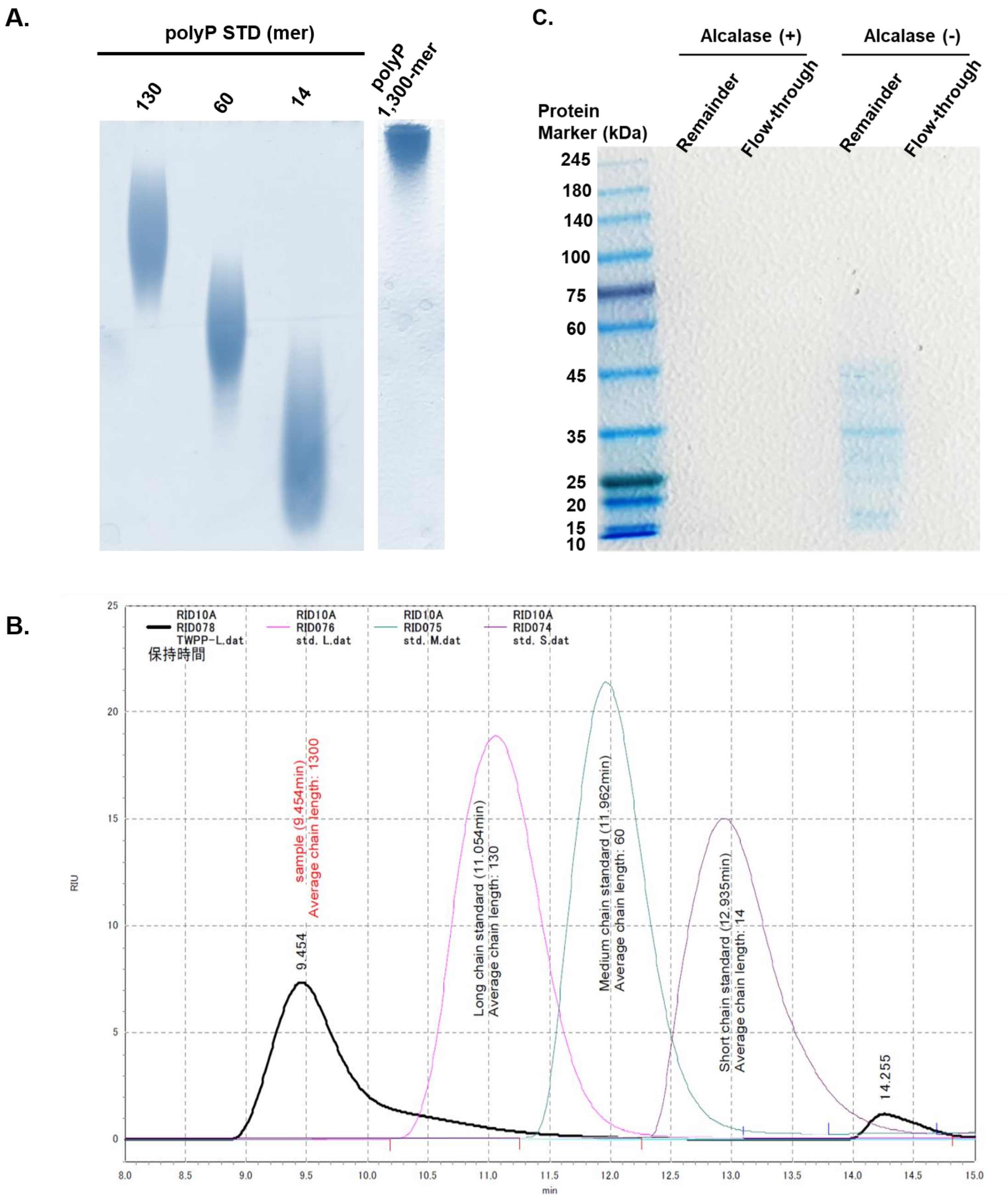


**Figure S6. Purification and determination of the length of the insoluble polyP product.** Related to Fig. 7. The reaction mixture containing the insoluble long-chain polyP was filtrated before and after protolysis treatment. **(A)** The insoluble long-chain polyP within the remainder fraction was then analyzed by TBE-Urea polyacrylamide gel electrophoresis, along with commercial polyP standards at various lengths (14-mer, 60-mer, 130-mer) as a standard to determine polyP product length. **(B)** The average length of the insoluble long-chain polyP was also determined *via* HPLC along with the same commercial polyP standards as standards to determine the product polyP length to be ~1,300-mer. **(C)** Removal of proteins from the remainder fraction following purification was verified by SDS-PAGE analysis.

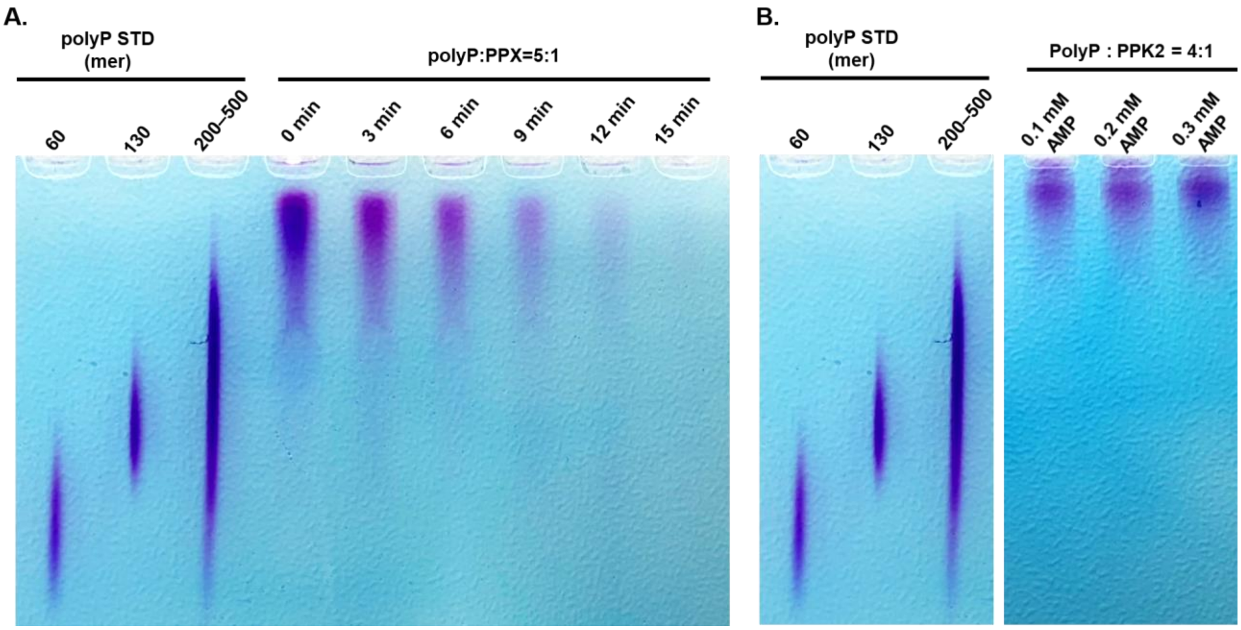


**Figure S7. Time-dependent enzymatic digestion of homogeneous long-chain polyphosphate (polyP) 1,300-mer using PPX or PPK2.** Related to Fig. 8. The polyP 1,300-mer was incubated with **(A)** PPX or **(B)** PPK2 to digest the polyP. The polyP and PPX or PPK2 were mixed at ratios of 5:1 or 4:1, respectively. The reaction mixtures collected at different time points were analyzed by TBE-Urea polyacrylamide gel electrophoresis, along with the commercial polyP standards of varying lengths as a reference for the lengths.

### Supplemental Tables

**Table S1. Complete list of recombinant proteins and expression vectors used in this work.**

| Enzyme | Abbreviation | Molecular weight (kDa) | UniProt accession | Host organism |
|---|---|---|---|---|
| ***Cytophaga* PPK2** | PPK2 | 72 | A0A6N4SMB5 | *Cytophaga hutchinsonii* |
| ***Ralstonia* PPK2c** | PPK2c | 42 | Q0KCB8 | *Ralstonia eutropha* |
| **Creatine kinase** | CK | 43 | P00567 | *Oryctolagus cuniculus* |
| **Exopolyphosphatase** | PPX | 58 | P0AFL6 | *Escherichia coli* |

**Table S2. $K_m$ and $k_{cat}$ of the enzymes used in this work (the values were extracted from the BRENDA database)**

| Enzyme | Substrate | $K_m$ (mM) | $k_{cat}$ (1/s) | $k_{cat}/K_m$ (1/s·mM) |
|---|---|---|---|---|
| ***Cytophaga* PPK2** | Polyphosphate | 0.3 ± 0.1 | 30.0 ± 5 | $1.0 \times 10^5$ |
| | AMP | 0.6 ± 0.10 | 17.0 ± 1 | $2.7 \times 10^4$ |
| | ADP | 0.4 ± 0.1 | 9.5 ± 0.5 | $2.7 \times 10^4$ |
| | ATP | 2.7 ± 0.7 | 0.3 ± 0.1 | $1.0 \times 10^2$ |
| ***Ralstonia* PPK2c** | ATP | 4.5 ± 2 | 10.0 ± 2 | $2.0 \times 10^3$ |
| **Creatine kinase** | ADP | 0.3 | N.A. | N.A. |
| | ATP | 0.7 | N.A. | N.A. |
| | Creatine | 5.0 | N.A. | N.A. |
| | Creatine phosphate | 0.8 | N.A. | N.A. |
| **Exopolyphosphatase** | N.A. | N.A. | N.A. | N.A. |

### Appendix

**SDS-PAGE gel images of purified *Cytophaga* class III PPK2 (~72 kDa), *Ralstonia* PPK2c (~42 kDa), rabbit creatine kinase (CK; ~43 kDa), and exopolyphosphatase (PPX; ~58 kDa).**

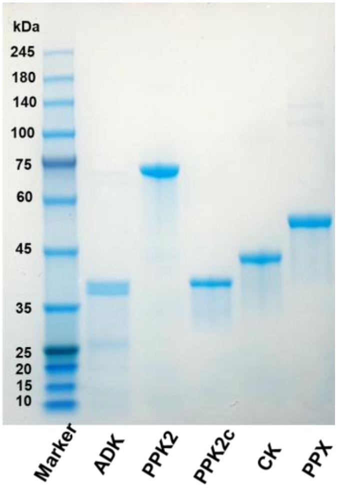


**HPLC chromatograms of commercial creatine phosphate (4.1 min), and creatine (10.0 min) standards.**

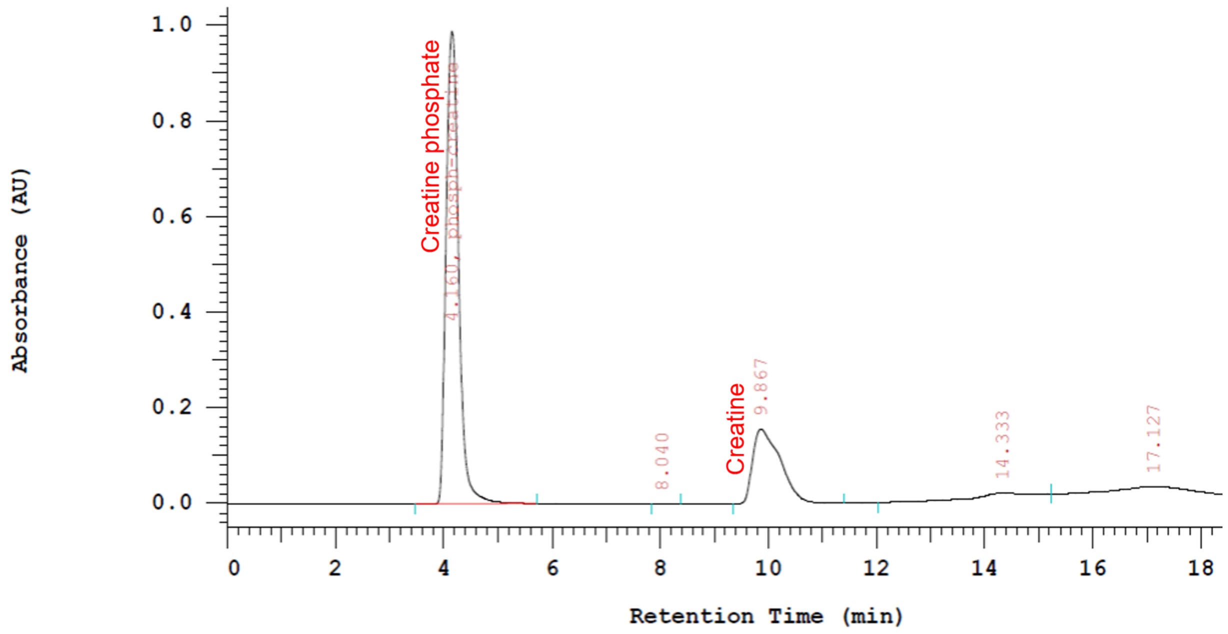


**Optical standard curves used in this study**

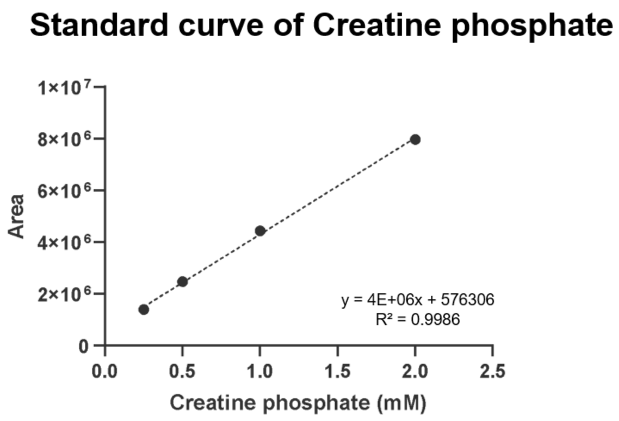


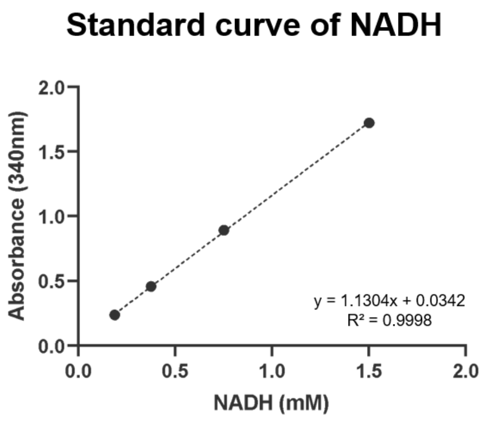


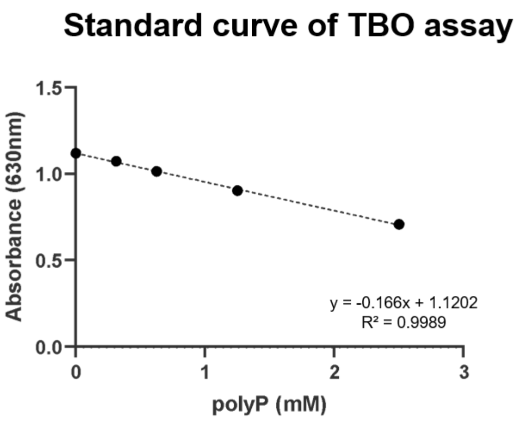

**The P-rich wastewater-sampling site**

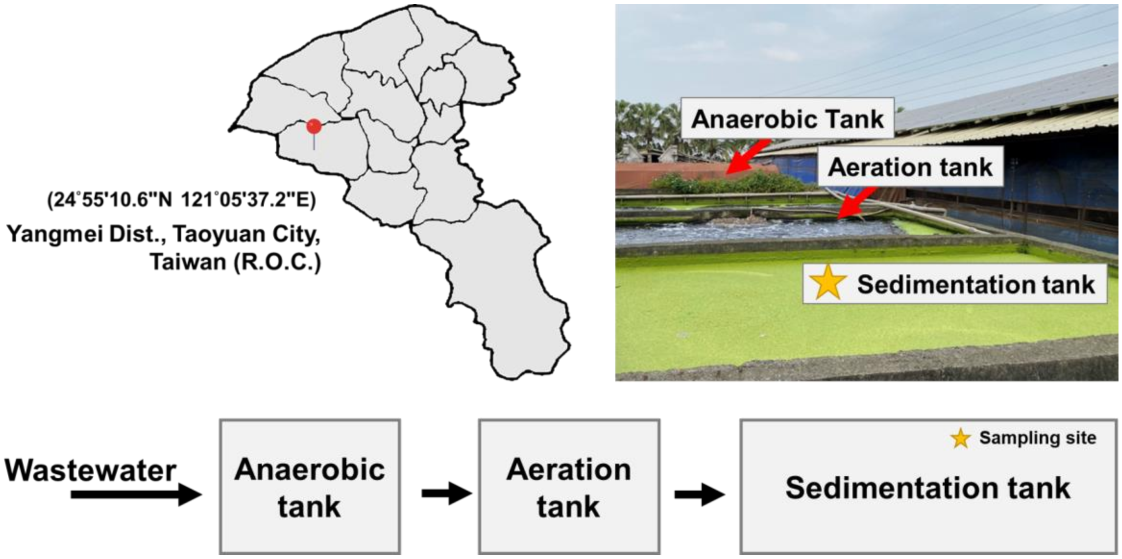
